# Quinone-transporting filaments extend the respiratory chain of Gram-positive bacteria

**DOI:** 10.64898/2026.06.26.734721

**Authors:** Ashleigh Kropp, Kazem Asadollahi, Jamie A. Stapleton, Luka Simsive, Pok Man Leung, Rachel L. Darnell, Christopher K. Barlow, Benjamin G. Hartmann, Nicholas D. J. Yates, Daniel R. Fox, Chris Greening, Oleksii Zdorevskyi, Vivek Sharma, James N. Blaza, Alison Parkin, Rhys Grinter

## Abstract

Cellular respiration depends on transferring electrons to hydrophobic quinones in membrane bilayers, constraining capacity to available surface area. Expanding this capacity is thought to have driven cellular complexity and eukaryogenesis, with Gram-negative bacteria evolving internal invaginations and eukaryotes using membrane-bound organelles. Whether Gram-positive bacteria, which lack such membranes, evolved alternatives was unknown. Here, we show that Bacillus subtilis forms a quinone-transporting pseudomembrane composed of filaments of the NADH dehydrogenase Ndh and the quinone-transporting protein Ncp. Cryo-EM, lipidomics, and molecular dynamics reveal that Ndh and Ncp co-assemble with phospholipids into a complex containing a solvent-excluded hydrophobic lumen that sequesters quinones. These complexes further assemble into filaments, linking chambers into a continuous conduit that amplifies quinone reduction while occupying minimal membrane space. Phylogenetic analysis suggests this recent innovation is widespread in Bacillota. Quinone-transporting filaments thus reveal a third strategy for overcoming surface-area limits and provide principles for engineering synthetic energy systems.

## Introduction

Cellular respiration is fundamentally constrained by the surface area available to house electron-transport machinery^1,2^. Overcoming this limitation has been proposed as a central driver of eukaryogenesis, with mitochondria and chloroplasts providing massively expanded internal membranes to support energy conversion^3–6^. Certain Gram-negative bacteria also employ organelles for energy conversion, notably thylakoids, chlorosomes, and anammoxosomes^7^. Others use extensive membrane invaginations to boost electron input capacity, as seen in methanotrophs, nitrifiers, and the presumed precursors of mitochondria^3,4,8–13^. Gram-positive bacteria appear to lack internal membranes or organelles that expand their respiratory capacity^14^. In addition, they only have a single membrane, which must accommodate other essential processes such as cell wall synthesis, nutrient import, and secretion^14,15^. Because respiratory ATP synthesis depends on electron transfer within this limited membrane surface, mechanisms that expand this capacity would confer an advantage under rapid growth or metabolic stress^1,16^. Central to this process are respiratory quinones, highly hydrophobic molecules that shuttle electrons between respiratory dehydrogenases and terminal oxidases^17,18^. For respiratory enzymes, accessing membrane-embedded quinones poses a fundamental challenge. Many enzymes solve this problem by using integral membrane subunits, containing heme or iron–sulphur clusters, that reduce quinones directly within the bilayer, and require permanent membrane real estate^19,20^. Others peripherally associate with the membrane to reach the quinone pool without dedicated transmembrane subunits^21,22^. Another recently discovered strategy involves extracting quinones from the membrane and reducing them in the cytosol using either membrane-associated quinone transport channels or a soluble quinone-binding shuttle protein^23–27^.

Nicotinamide adenine dinucleotide (NADH) is a key intermediate electron carrier in cellular metabolism, transporting high-energy electrons primarily from organic carbon oxidation to the electron transport chain via the reduction of respiratory quinone^28,29^. While this reaction is classically associated with the multi-subunit complex I (type-1 NADH dehydrogenase), many bacteria instead rely on type-2 NADH dehydrogenase (NDH-2). Unlike complex I, NDH-2 is a compact, ∼50 kDa flavoprotein that does not span the membrane or pump protons^30–34^. NDH-2 performs catalysis using a single flavin redox cofactor, with NADH binding to one side and quinone to the other. While NDH-2 does not conserve energy via proton pumping, it is often the dominant or sole NADH-oxidising enzyme in bacteria, underscoring its physiological importance^21,35^. Its possible advantages are twofold. First, the large midpoint reduction potential (Δ*E*_m_) gap between NADH (–320 mV) and quinones (menaquinone, –70 mV; ubiquinone, +60 mV) makes NDH-2 catalysis essentially thermodynamically irreversible, ensuring robust forward electron flux even under a strong proton-motive force that might reverse complex I^32^. Second, the small size and low biosynthetic cost of NDH-2 make it well suited to conditions where resource availability is uncertain, membrane or cytosolic real estate is limited, and/or rapid growth is prioritised. NDH-2 exemplifies a minimalist yet highly effective solution for sustaining electron flow across diverse microbes and lifestyles^36^.

The soil-dwelling facultatively anaerobic Gram-positive bacterium *Bacillus subtilis* encodes three putative NDH-2 dehydrogenases, *ndh, yumB and yutJ,* to transfer electrons from NAD(P)H to the respiratory chain^35,37^. Among these, Ndh is the primary NADH dehydrogenase during vegetative growth^35^. NDH-2 enzymes often peripherally associate with the membrane to transfer electrons to quinone within the bilayer ^21,32,38–42^. However, *B. subtilis* Ndh lacks key residues in this region for membrane interaction, raising the question of how it accesses and reduces quinones. The *ndh* gene shares an operon with *yjlC*, which encodes a protein with a C-terminal DUF1641 domain^35^. Recently discovered DUF1641-domain proteins form quinone-transporting complexes with both H₂- and formate-oxidising enzymes (Huc and ForCE) ^23,24^. Notably, the DUF1641 protein HucM from *Mycobacterium smegmatis* assembles into a central tetrameric tube that scaffolds eight HucSL subcomplexes. The resulting Huc complex contains an internal hydrophobic chamber that connects to the plasma membrane via the HucM tube, enabling quinone extraction and its reduction within the complex^24^.

Here, we identify that the *yjlC* gene product, which we designate the Ndh-coupling protein (Ncp), forms a stable NADH-oxidising complex with Ndh. Unlike the Huc and ForCE complexes, the Ndh-Ncp complex assembles into tube-like filaments with internally connected hydrophobic chambers lined by many embedded phospholipids^23,24^. These Ndh-Ncp ‘pseudomembrane’ tubes enable the sequestration and long-range transport of respiratory quinones, hundreds of angstroms outside of the membrane, for reduction by electrons from NADH. By doing this, Ndh–Ncp filaments extend the surface area for quinone reduction, providing a solution to the limitation of membrane real estate available for energy generation, functionally convergent with mitochondrial cristae and respiratory super complexes^3–6^. Genomic analysis shows that Ndh–Ncp complexes are widely distributed across Bacillota, indicating that respiratory chain expansion via protein-lipid pseudomembrane filaments is a conserved and fundamental mechanism in bacterial bioenergetics.

## Results

### Ndh from *B. subtilis* forms a functional complex with Ncp

The *ndh* gene from *B. subtilis* is important for optimal growth and forms an operon with *ncp* (formerly *yjlC*)^35^. To test whether *ndh* and *ncp* are functionally associated, we generated *B. subtilis* knockout strains lacking either gene (Δ*ndh*::kan or Δ*ncp*::kan) and assessed their growth in liquid culture. Both Δ*ndh*::kan and Δ*ncp*::kan strains exhibited a slower growth rate, with defects of similar magnitude at 37°C, suggesting that loss of *ncp* phenocopies loss of *ndh* (**Figure 1a; Figure S1a**). In addition, when grown at 30°C (which reduces sporulation of *B. subtilis*), the growth yield of both strains was significantly reduced (**Figure S1b,c**).

**Figure 1.**
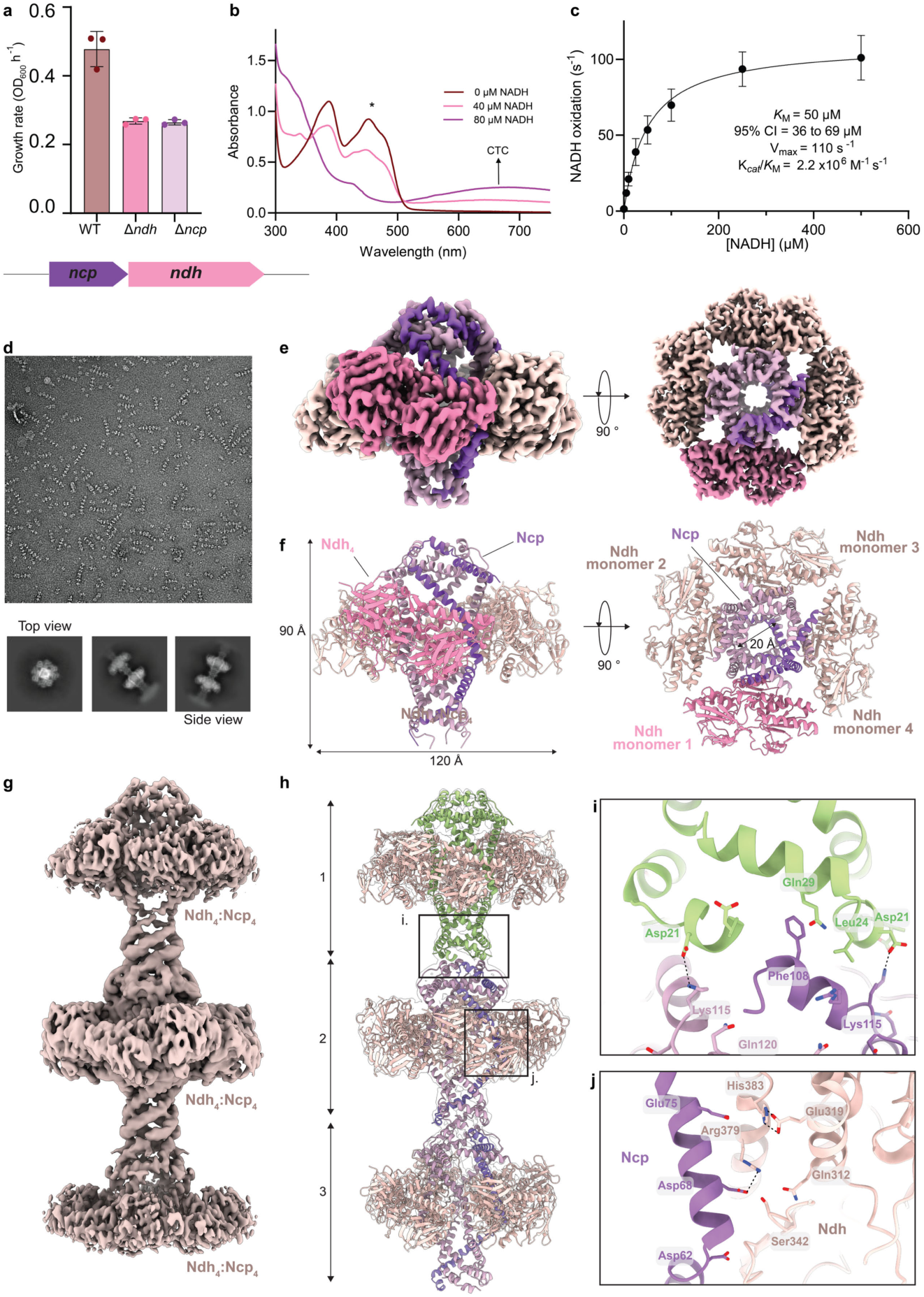
*B. subtilis* Ncp complexes with Ndh and is essential for its activity. a, Growth rates of WT, Δ*ndh* or Δ*ncp B. subtilis* 168 strains during exponential phase. Data presented is from three biological replicates (n = 3) with mean ± standard deviation displayed. **b,** Representative UV-visible spectra of Ndh-Ncp complex with increasing concentrations of NADH. The asterisk (*) shows the typical flavin peak at 450 nm, and the arrow represents the charge-transfer-complex (CTC) at ∼660 nm. **c,** Michaelis-Menton kinetics of Ndh-Ncp incubated with MDA and varying concentrations of NADH. Data presented is from three independent replicates (n = 3) with mean ± standard deviation displayed. **d,** A representative negative stain EM micrograph showing filamentous Ndh-Ncp (above) and cryo-EM 2D class averages of filaments (below). **e,** A cryo-EM map of the Ndh-Ncp complex showing the Ndh subunits in pale pink and one representative in bright pink, and the Ncp in purple. The left image shows the side view, and the right image shows the top view of the complex. **f,** A cartoon representation of the cryo-EM structure of Ndh-Ncp complex. Four Ndh monomers (light pink) couple to Ncp (purple) with C4 symmetry. The left image shows the side view, and the right image shows the top view of the complex. **g,** A cryo-EM map of the Ndh-Ncp helical refinement showing three subunits in the filament**. h,** The Ndh-Ncp filament structure showing three units in the filament. A cartoon representation is shown with the top Ncp tube in green and the following two tubes in purple and the interacting Ndh enzymes in pale pink. **i,** The interaction between the ‘tail’ (green) of Ncp from one unit and the ‘head’ (purple) Ncp from another unit in the filament. Key residues that promote filament formation are highlighted. **j,** The interface between one Ndh enzyme and Ncp monomer, highlighting key residues that drive the interaction.

Given the genetic linkage between *ndh* and *ncp*, and the homology of Ncp to HucM, we hypothesised that Ndh and Ncp form a complex that couples Ndh activity to the membrane. To investigate this, we constructed an *Escherichia coli* expression construct encoding both *ncp* and *ndh* genes in tandem, with *ncp* N-terminally twin-strep tagged. Expression and purification from clarified cell lysate resulted in the isolation of a complex containing both Ndh and Ncp, which had an elongated peak on size exclusion chromatography (SEC), indicating a non-uniform size distribution (**Figure S1d**). UV-visible spectra demonstrated that the purified Ndh-Ncp complex contains flavin in an oxidised state with absorbance peaks at ∼390 nm and 450 nm. The addition of NADH led to the formation of a charge-transfer complex (CTC) between NADH and FAD⁺, indicated by an absorbance peak at 660 nm (**Figure 1b, Figure S1e**)^32^. This CTC wavelength is comparable to that of NDH-2 from *Caldalkalibacillus thermarum* and *Staphylococcus aureus* but is shorter than the 740 nm of the distantly related eukaryotic NDH-2 from *Saccharomyces cerevisiae*^32,40,43^. This implies that the mechanism of NADH oxidation by Ndh is similar or identical to that of other bacterial NDH-2 enzymes^32^.

Next, we examined whether the Ndh-Ncp complex displays catalytic kinetics comparable to other NDH-2 homologues. Because other NDH-2 enzymes directly reduce respiratory quinone, we utilised menadione (MDA, the redox active headgroup of menaquinone) or decylubiquinone (DQ) as the electron acceptor for these experiments. Ndh-Ncp oxidises NADH with a *K*_M_ of 50 µM (CI 36 to 69 µM) and a V_max_ of 110 s^-1^, with MDA as the electron acceptor (**Figure 1c; Figure S1f**). Ndh-Ncp has lower affinity and is ∼10-fold slower than NDH-2 from *C. thermarum* with MDA as an electron acceptor (K_M_ = 29 µM, V_max_ =1190 s^-1^) ^32^. Using DQ as an electron acceptor for Ndh-Ncp resulted in a lower V_max_, suggesting that Ndh-Ncp prefers menaquinone over ubiquinone (**Figure S1g,h**). The slower kinetics of *B. subtilis* Ndh-Ncp compared to *C. thermarum* NDH-2 suggest this complex may restrict the access of soluble quinones to the Ndh electron acceptor sites compared to characterised NDH-2 enzymes.

### Ndh and Ncp form a complex that associates into filaments

To gain insight into the structure of the Ndh-Ncp complex, we performed negative stain electron microscopy. Strikingly, these micrographs revealed that purified Ndh-Ncp forms filaments with numerous repeating units. To determine the high-resolution structure of this complex, we performed cryo-electron microscopy (cryo-EM) on Ndh-Ncp, followed by single particle and helical reconstruction (**Figure 1d**). We resolved the structure of a single Ndh-Ncp complex at a global resolution of 2.11 Å, revealing a 4:4 stoichiometry of Ndh and Ncp subunits (**Figure S2, Supplemental Table S1**). Ncp forms a hollow tubular tetrameric tube at the centre of the complex, which scaffolds four Ndh subunits. The complex has C4 symmetry, spans ∼90 Å in length and ∼120 Å in width, and has openings in the Ncp tube of ∼20 Å in diameter at its head and tail (**Figure 1e,f**). The interface between the Ndh monomers and the Ncp scaffold is primarily driven by electrostatic interactions mediated by a positively charged Ndh C-terminal helix, which is complemented by the negative charge of the Ncp scaffold in this region (**Figure 1j, Figure S1i**). To reconstruct the Ndh-Ncp filament, we performed helical refinement, generating a map with three connected Ndh-Ncp complexes (**Figure 1g,h, Figure S2, Supplemental Movie S1, Table S1, Figure S1j**). These Ndh-Ncp subunits stack head-to-tail to form the filament, with the connecting interface composed entirely by the Ncp subunit and driven by a phenylalanine (Phe108) on the Ncp head embedding into the tail complex, as well as a salt bridge between Lys115 (head) and Asp21 (tail) (**Figure 1h,i**).

The width of the Ncp tube varies, reaching ∼45 Å in the region where Ndh attaches (**Figure 2a**). The C-terminal ‘head’ of the Ncp tetramer contains the DUF1641 domain. This region is structurally similar to HucM^24^, with each Ncp subunit travelling up and over the monomer immediately left of it, then under the monomer immediately right of it, stabilising the complex and creating the opening at the head of the tube (**Figure 2b,c**). In Huc, this region is responsible for interaction with the cell membrane, containing positively charged amino acids, which mediate interaction with phospholipid head groups, and a tryptophan that stabilises interactions with the bilayer^24^. Consistent with this function, this region of the Ncp tetramer is positively charged, with Phe108 positioned to embed into the hydrophobic region of the lipid bilayer (**Figure 2c,d**). Interestingly, this region is also responsible for filament formation, suggesting a dual role in lipid and protein interactions, reminiscent of the recently characterised CODH-CoxG complex^44^. Considering Ndh-Ncp forms filaments and that the complex is purified from the soluble fraction, interaction between the Ncp and the membrane is most likely transient.

**Figure 2.**
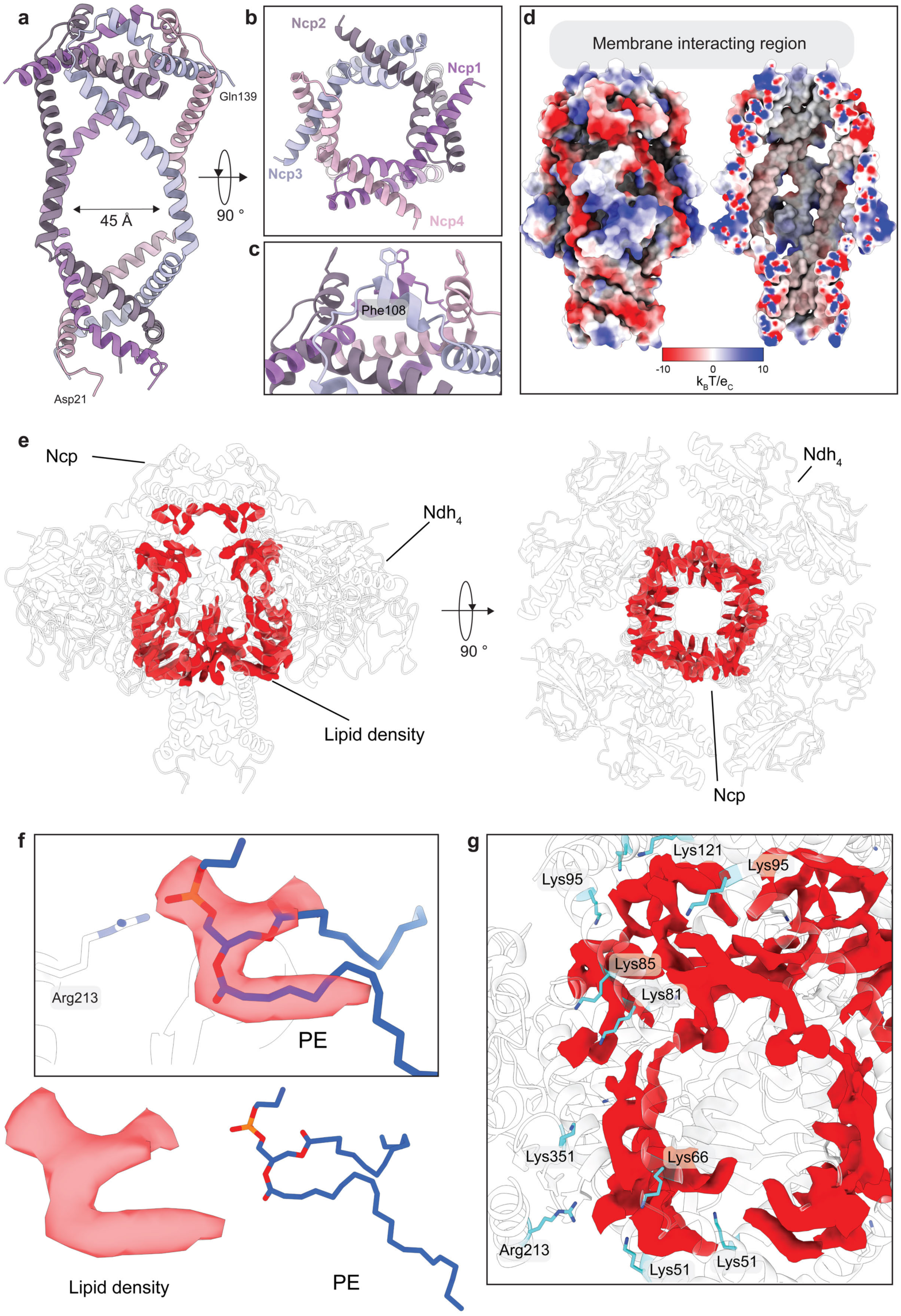
Ncp facilitates membrane interaction, and Ndh-Ncp associates with lipids. a, A cartoon representation of the structure of Ncp tetramer alone, with each Ncp monomer in a di_erent shade of purple. The Ncp tube broadens where Ndh binds, reaching a width of ∼45 Å. **b,** The DUF1641 domain forms the opening of the tube and **c,** Phe108 from each monomer facilitates membrane interaction. **d,** The membrane interacting region of Ncp is characterised by positively charged residues (blue) flanking the hydrophobic Phe108 (white) that inserts into the hydrophobic bilayer. Internally, the Ndh-Ncp complex is hydrophobic (white). **e,** A cartoon representation of the Ndh-Ncp cryo-EM structure in white with black outline, with density corresponding to phospholipids in red. **f,** An example of a phosphatidylethanolamine (PE) molecule modelled into lipid density from the cryo-EM maps of the Ndh-Ncp complex, with the phosphate head group coordinated by Arg213, **g,** The positively charged residues in blue from the Ndh and Ncp subunits that coordinate lipid density (red) throughout the complex.

The Ndh-Ncp filament consists of a tube formed by the head and tail region of the Ncp tetramer, which connects a broader chamber where Ndh associates with the complex. Both the tube and chamber are lined entirely with hydrophobic amino acids, forming a continuous conduit for hydrophobic quinones from the point of membrane attachment to the electron acceptor sites of Ndh (**Figure 2d**). Strikingly, additional density is present in the cryo-EM maps in the region of Ndh attachment, filling gaps between the Ndh and Ncp polypeptide chains (**Figure 2e, Supplemental Movie S2**). The shape and chemical environment of this density are consistent with it belonging to phospholipids. The solvent facing portion of these densities, which likely corresponds to negatively charged phosphate groups of phosphatidylglycerol (PG) and phosphatidylethanolamine (PE) lipids, is coordinated by complementary positively charged residues (Lysines 51, 66, 81, 85, 95, and 121 from Ncp and Arg213 and Lys351 from Ndh), and the density likely corresponding to aliphatic tails extends into the hydrophobic interior of the Ndh-Ncp complex (**Figure 2e,f,g**). The presence of this density indicates that the Ndh-Ncp complex incorporates a significant quantity of phospholipids as an additional structural and functional element. Overall, the structure of the Ndh-Ncp complex reveals similarities to Huc and ForCE, indicating a conserved role for the DUF1641 superfamily in enzyme scaffolding, membrane attachment and long-range quinone transport^23,24^. However, filament formation by Ndh-Ncp and the incorporation of phospholipids into the structure indicate significant structural and mechanistic differences and demonstrate pronounced structural diversity within the DUF1641 superfamily.

### Ndh-Ncp binds NADH and menaquinone in conserved binding sites

Our initial cryo-EM data were collected on Ndh-Ncp in the absence of NADH. This structure had no NADH or quinone bound at their respective binding sites. To obtain the substrate-bound structure, we repeated the cryo-EM, supplementing Ndh-Ncp with NADH immediately before freezing. We resolved this complex at a global resolution of 2.52 Å, showing clear density for the FAD group, NADH and quinone, at their structurally conserved binding sites (**Figure 3a-e, Figure S3, Supplemental Movie S3, Supplemental Table S1**). FAD coordination by Ndh is comparable to other structurally characterised NDH-2 enzymes, with isoalloxazine stabilisation from Lys127, Glu168 and Lys373 side-chain residues and Ala311 and Gln312 main-chain nitrogens^32,34,38–41^. The pyrophosphate groups engage in hydrogen bonding with the Tyr12, Gly13 and Asp295 main-chain nitrogens, and the adenine ring interacts with the Val80, Gln261, and Lys37 main-chain nitrogens (**Figure 3b,c**)^32,38–41^. Ndh binds NADH in the conserved region, interacting with the key WxxG motif (Trp254 and Gly257), as well as interactions with the highly conserved residues Ile116, Gly163, Val196-Gly199, Val231, Pro309 and Thr310 (**Figure 3d**)^32,38–41^.

**Figure 3.**
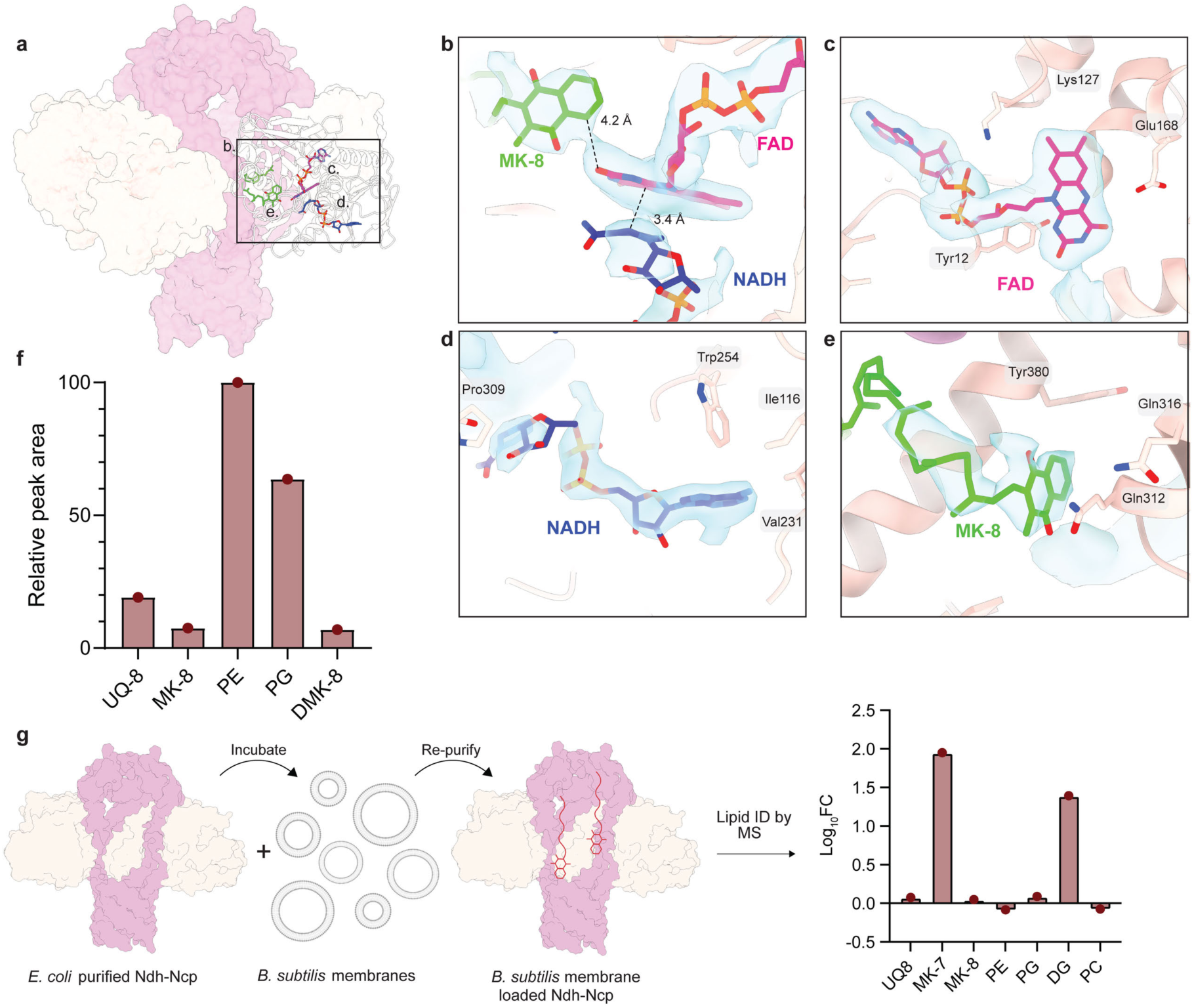
Ndh-Ncp binds NADH and MK-8. a, Cartoon representation of the cryo-EM structure of Ndh-Ncp with NADH and quinone bound, showing their location in one Ndh enzyme. **b,** The interatomic distances between each of the ligands, NADH in blue, FAD in pink and MK-8 in green. **c,** The coordination of the FAD is driven by conserved Lys127, Glu168, Lys373 and Tyr12. **d,** NADH has a conserved binding mode. **e,** MK-8 is coordinated by Tyr380, Ala311, Gln312 and Gln316. Density for all three cofactors is shown overlayed on the structure, contoured at 5σ. **f,** Graph showing the relative peak area of lipids in the Ndh-Ncp purified sample, normalised to the most abundant lipid, PE. Data were from positive ion mode MS and each type of lipid was summed into its associated group (i.e., PE lipids of di_ering lengths were summed into the PE group). **g,** Schematic of the experimental protocol for the incubation of Ndh-Ncp with *B. subtilis* membranes and graph showing the fold change in lipids and quinones after treatment. PE = phosphatidylethanolamine, PG = phosphatidylglycerol, UQ-8 = ubiquinone-8, MK-8 = menanquinone-8, MK-7 = menaquinone-7, DMK-8 = demethylmenaquinone-8, DG = diglyceride, PC = phosphatidylcholine. Lipidomics data in panels **f** and **g**, is derived from a single experiment (n = 1).

In our NADH-bound structure, we observed density at the conserved quinone-binding electron acceptor site^38,39,41^. The absence of this density in the NADH-free structure may indicate that quinone binding is preceded by NADH binding, leading to the formation of the CTC observed in our spectroscopic analysis (**Figure 1b, Figure S1e**). *E. coli*, the expression host for the Ndh-Ncp complex, produces ubiquinone-8 (UQ-8) and menaquinone-8 (MK-8) as its major quinones during aerobic and anaerobic conditions, respectively^17^. The shape and coordination environment of the quinone headgroup density in our structure is consistent with MK-8, so we modelled this in our structure (**Figure 3e, Figure S4a,b**). This is also supported by the superior kinetics for Ndh-Ncp with MDA versus DQ as the electron acceptor (**Figure S1f,h)**^45^. The Ndh quinone binding site is conserved with other NDH-2 homologues, specifically the motif (AQxAxQ; Ala311, Gln312, Ala314, Gln316)^32,38,41^. The MK-8 binding site faces the interior of the Ndh-Ncp tube, indicating that quinone enters the Ndh electron acceptor site from the internal hydrophobic chamber of the complex (**Figure 3e, Figure S4c,d**). Considering this, the slower kinetics of Ndh-Ncp are likely to result from restricted access to the electron acceptor site, due to the tube slowing the movement of the soluble quinones used in this assay (**Figure 1c**).

### The Ndh-Ncp complex binds phospholipids and extracts menaquinone

Our cryo-EM structure suggests that phospholipids form an important component of the Ndh-Ncp complex, and that the hydrophobic chamber of the complex contains respiratory quinones. To identify lipids associated with the complex, we performed lipidomics on the purified Ndh-Ncp complex. This analysis revealed considerable quantities of lipids associated with the complex, with the greatest combined peak area corresponding to PE and PG lipid species (**Figure 3f, Supplemental Data S1**). This validates our interpretation of the density in our cryo-EM structure as PE and PG phospholipids. These lipids are most likely derived from the *E. coli* plasma membrane and associate with the complex during or after assembly. The quantity of lipids associated with the complex tube is remarkable and illustrates a novel mechanism of membrane extension through a protein-lipid hybrid complex.

In addition to phospholipids, abundant features with masses and elution profiles corresponding to ammonium adducts of UQ-8, MK-8 and DMK-8 were associated with the complex (**Figure 3f, Supplemental Data S1**). The detection of MK-8 supports its modelling at the electron-acceptor site in our NADH-bound structure. In addition, UQ-8 represents the largest individual peak area in our lipidomics data, and the total peak area ratio of quinones to phospholipids is ∼1:5 (**Figure 3f**). While the detection efficiency of different lipids can vary considerably in LC-MS analysis, these data indicate that quinones are abundant in the Ndh-Ncp complex. A mass balance estimate yields an expected quinone to phospholipid ratio ∼1:100–1:300 in *E. coli* (See methods), indicating the Ndh-Ncp complex is concentrating quinones relative to the membrane. The presence of both ubiquinone and menaquinone suggests this sequestration is driven primarily by interaction with their hydrophobic isoprenyl tails, which would be readily accommodated by the hydrophobic interior of the complex. Given the high quinone content, it is likely that quinones occupy both canonical Ndh quinone binding sites and the central Ndh-Ncp cavity, although direct structural assignment of additional quinones was not possible due to the dynamic nature of this hydrophobic environment.

To assess whether the Ndh-Ncp complex can extract quinones from native membranes, we incubated the purified complex with isolated *B. subtilis* membranes, then re-purified the complex and analysed its lipid content. *B. subtilis* produces MK-7 as its sole respiratory quinone, and accordingly, MK-7 was the only quinone species with increased abundance^45^. Following incubation, MK-7 levels in the Ndh-Ncp sample increased by ∼90-fold relative to pre-incubation (**Figure 3g, Figure S4e,f, Supplemental Data S1**), indicating substantial quinone uptake. The sequestration of quinones by the Ndh-Ncp complex could allow quinone reduction when access to the membrane-bound quinone pool is limited, allowing more respiratory complexes to operate in parallel, and increasing the effective surface area of the respiratory chain.

### The architecture of the Ndh-Ncp-phospholipid complex

Our structural and lipidomic analysis indicates that Ndh-Ncp forms a hybrid protein-phospholipid complex. However, the density in our cryo-EM maps is insufficiently resolved to model the location of phospholipids in this complex. To resolve this, we performed molecular dynamics (MD) simulations of Ndh-Ncp in complex with PE and PG and associated with a PE/PG lipid bilayer. Due to the size and complexity of the system, we performed extensive optimisation simulations to determine optimal lipid numbers and starting positions (**Supplemental Note S1, Figure S5a, Supplemental Table S3**). The Ndh-Ncp chamber is significantly larger than the Huc or ForCE complexes^23,24^, allowing it to accommodate this comparatively large number of phospholipids, 28 PE and 7 PG in total. In this model, the charged phosphate-containing head groups of PE and PG occupy gaps in the complex between the Ndh and Ncp subunits, facing towards the external solvated environment (**Figure 4a,c,d**). The phospholipid headgroup positioning and coordination correlate well with our cryo-EM data, indicating the MD simulations provide a realistic model of the complex (**Figure 4b**). The aliphatic lipid tails extend into and occupy the internal chamber of the complex (**Figure 4c,d**). The lipid tails are dynamic throughout the simulations, which is consistent with the poor density in our cryo-EM maps and the role of the complex in long-range quinone transport, which requires lipid mobility within the chamber.

**Figure 4.**
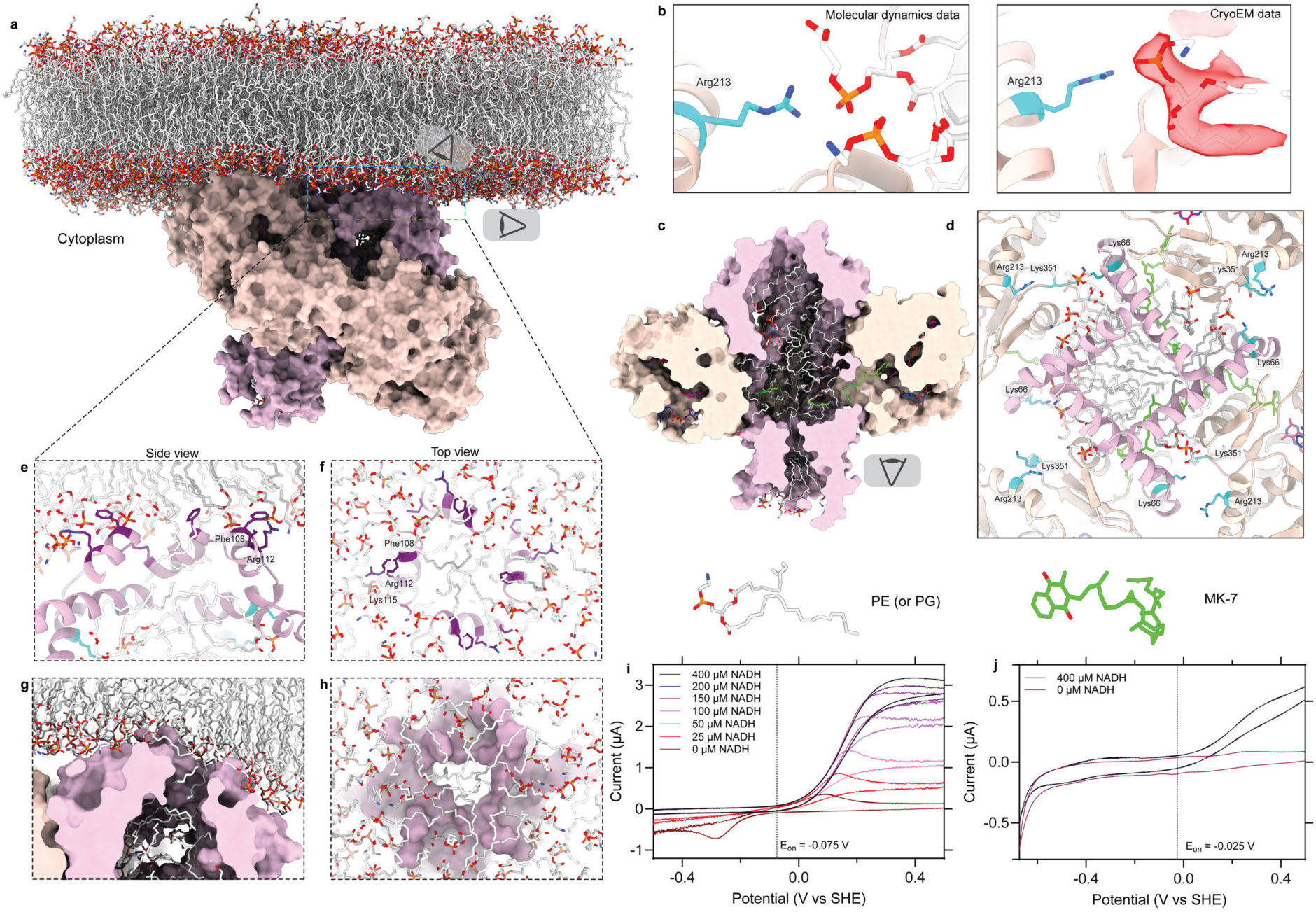
Molecular dynamics simulations demonstrate the Ndh-Ncp protein-lipid architecture a, The Ndh-Ncp simulation was setup with a bilayer consisting of PE and PG lipids. Lipids and MK-7 were docked inside the Ncp based on structural and lipidomics data. **b,** A PE and PG molecule from a MD snapshot showing the PG coordinated by Arg213, left, and cryo-EM data showing a PE molecule in the cryo-EM density coordinated by Arg213, right. **c,** Snapshot of the last frame of the simulation with PE/PG present within Ndh-Ncp. Cut away shows the occupation of these molecules within the Ncp and quinone binding site. MK-7 is shown in green and PE/PG in white. **d,** Snapshot of the last frame of the simulation showing the position of phospholipid headgroups in proximity to positively charged residues predicted to coordinate the lipid headgroups (lipids in white, coordinating residues in cyan). The complex is viewed from the bottom. **e,** Side view and **f,** top view snapshot from the last frame of the MD simulation of the Ncp interacting with the bilayer. Key residues Phe108, Arg112 and Lys115 are shown in dark purple. **g,** Side view and **h,** top view of the last frame from MD data of the Ncp tube interacting with the bilayer. Lipids from the bilayer enter the Ncp with the hydrophobic tail directed into the Ncp tube. **i,** Representative cyclic voltammograms of Ndh-Ncp stabilised on a phospholipid membrane recorded at varying NADH concentrations from-0.675 to +0.625 V vs SHE at 10 mVs^-1^ in the presence of MK-7. **j,** Representative cyclic voltammograms of Ndh-Ncp recorded at 0 µM NADH (pink) or 400 µM NADH (black) from-0.675 to +0.625 V vs SHE at 10 mVs^-1^ in the absence of exogenous MK-7.

As indicated by our structural analysis, the Ncp tube interacts with the membrane through four Phe108 residues that stably embed in the bilayer and four Arg112 and Lys115 residues that coordinate the charged phospholipid headgroups (**Figure 4e,f**). Over the course of the simulations Ndh-Ncp complex tilts so that one of the Ndh subunits contacts the membrane, with the opening of the Ncp tube penetrating the hydrophobic region of the lipid bilayer (**Figure 4a,g, Figure S5b**). To accommodate the Ncp tube in the membrane, the phospholipid headgroups of the bilayer move away from the opening of the Ncp tube, and tails of bilayer phospholipids occupy the opening of the Ncp tube (**Figure 4g,h**). This arrangement connects the hydrophobic lumen of the bilayer with the interior of the Ncp-Ndh complex, allowing for diffusion of quinone between the two structures, a process critical for long-range quinone transport. This model contrasts with the recently published cryo-EM structure of the ForCE complex^23^. The architecture of ForCE is similar to the Huc and Ndh-Ncp complexes, with a tetramer of ForE subunits forming a membrane-associated quinone transport tube^23,24^. This structure contains phospholipid headgroups modelled within the hydrophobic ForE tube, with the hydrophobic tails directed towards the surrounding bilayer. However, the assignment of these phospholipid headgroups is not strongly supported by the cryo-EM density. This arrangement appears difficult to reconcile with the hydrophobic environment of the tube and would occlude the proposed quinone access pathway to the ForCE chamber, suggesting that this model may not accurately represent the architecture of this region^23^. Our MD data may generate a more chemically realistic model for Ndh-Ncp and DUF1641 domain interaction with the membrane.

### Quinone diLusion drives electron transfer from Ndh-Ncp to the membrane

To directly assess the ability of the Ndh-Ncp complex to utilise quinone to transfer electrons from NADH oxidation to a lipid bilayer, we performed cyclic voltammetry on the complex immobilised on an electrode-bound, MK-7-supplemented phospholipid membrane, stabilised by a mixed self-assembled monolayer consisting of EO_3_-cholesterol and 6-mercaptohexanol. In the absence of NADH, MK-7 oxidation and reduction peaks were observed in this system at +0.09 V and −0.28 V vs. SHE, respectively (**Figure 4i**). The addition of NADH yielded a catalytic oxidative current with an onset potential of-0.075 V, close to the midpoint potential of MK-7^46^, indicating that MK-7 is responsible for mediating electron transfer between the electrode and the Ndh. The reductive MK-7 peak disappeared upon NADH addition, indicating that MK-7 is reduced at the Ndh active site rather than at the electrode. The electrochemical *K*_M_ for NADH, determined from the magnitude of the catalytic current at +0.3 V vs. SHE at different NADH concentrations, was 91.6 µM, in agreement with our prior kinetic assays (**Figure S6a**). Control experiments lacking the enzyme showed no NADH oxidation (**Figure S6b**). These observations are consistent with the behaviour of other NDH-2 enzymes, including homodimeric *C. thermarum* NDH-2^47^.

We next examined the electrochemical behaviour of the complex in the absence of MK-7 in the electrode-associated bilayer. Cyclic voltammetry revealed that under these conditions, Ndh-Ncp produced a catalytic oxidative current upon the addition of 400 µM NADH, but at a reduced magnitude. We attribute this activity to the presence of MK-8 in Ndh-Ncp sequestered from *E. coli,* which decayed over repeated scans, suggesting that quinones present in the Ndh-Ncp complex are responsible for electron transfer from NADH to the electrode, and the magnitude of this transfer reduces over time as they diffuse into the lipid bilayer (**Figure 4j, Figure S6c**). We calculated a quinone diffusion constant of 8.2 × 10⁻⁴ s⁻¹ from the decay rate (**Figure S6c**), indicating the rate of diffusion between Ndh-Ncp and the membrane. Together with our lipidomics analysis showing substantial quinone enrichment by Ndh-Ncp, these electrochemical data demonstrate that the complex not only sequesters quinones from the membrane but also retains them in quantities capable of sustaining Ndh catalysis away from the bilayer, providing a mechanism for extending respiratory activity beyond the membrane.

Cyclic voltammetry experiments of Ndh-Ncp immobilised on a bilayer containing UQ-8 demonstrated Ndh is selective for MK-7, with a much greater onset potential (>0.5 V) observed for UQ-8 (**Figure 4j, Figure S6c,e**). This is consistent with kinetics experiments and further indicates that the MK-8 associated with the Ndh-Ncp complex is responsible for facilitating electron transfer in the electrochemical experiments in the absence of quinone supplementation in the bilayer.

### The Ndh-Ncp complex is widespread throughout Bacillota

To understand the distribution and evolution of Ndh-Ncp in Gram-negative bacteria, we performed a comprehensive survey of Ndh and Ncp homologues in all 113,104 representative archaeal and bacterial species genomes from the Genome Taxonomy Database (GTDB) (**Supplemental Data S2**)^48^. Ndh-Ncp complex is exclusively found in the phylum Bacillota, where 8.5% of genes encoding Ndh homologues occur directly adjacent to a *ncp* gene. The 684 Ndh-Ncp encoding species span 30 different families, indicating that this complex is widely encoded by taxonomically diverse members of this phylum (**Figure S7a**). Genome-wide metabolic analysis suggested that the Ndh-Ncp complex occurs in aerobic or facultatively anaerobic lineages but is absent in obligately anaerobic lineages such as Clostridia and Negativicutes (**Supplemental Data S2**). All Ndh-Ncp encoding species co-encode at least one additional, non-Ncp-associated NDH-2 enzyme (**Figure S7a**), suggesting distinct physiological roles of these enzymes, as exemplified by *yumB* and *yutJ* encoded by *B. subtilis*^35^.

Phylogenetic analysis of Ndh homologues across Bacillota revealed three major monophyletic NDH-2 clades (A, B, C) and three smaller clades (D, E, F) (**Figure 5a,b, Figure S8a, Figure S9f, Supplemental Data S2**). All Ncp-associated NDH-2 enzymes are contained within clade C where Ncp-associated Ndh enzymes form a monophyletic group, which we refer to as Clade C1, defined by *ndh–ncp* synteny and the evolution of an Ndh C-terminal domain that mediates the stable interaction interface with Ncp (**Figure 5a, Figure S7a, S9e**). In addition, we identify a transitional lineage that we term clade C2. Enzymes in clade C2 retain a C-terminus widely conserved in the NDH-2 family that mediates direct membrane interaction (**Figure 5a,b, Figure S8**). Phylogenetic analysis resolves two C2 sub-branches distinguished by the presence or absence of a DUF1641-containing gene within their genome (**Figure 5a, Figure S8b**). The DUF1641-encoding genes of this C2 branch lack the strict *ndh–ncp* synteny that defines clade C1, suggesting an intermediate stage in which the *ncp* progenitor was acquired by the ancestral bacterium, possibly forming a complex with a different enzyme, but the NDH-2 enzyme continued to interact with the membrane directly. From this transitional state, we hypothesise that the Ncp-associated Ndh enzymes in clade C1 emerged via spontaneous complex formation with the DUF1641-containing protein, followed by co-evolution to form the Ndh-Ncp. The short branch length of clade C1 relative to other clades suggests the complex formation with Ncp is a relatively recent evolutionary innovation, but has become widely disseminated in Bacillota (**Figure 5a, Figure S8a**).

**Figure 5.**
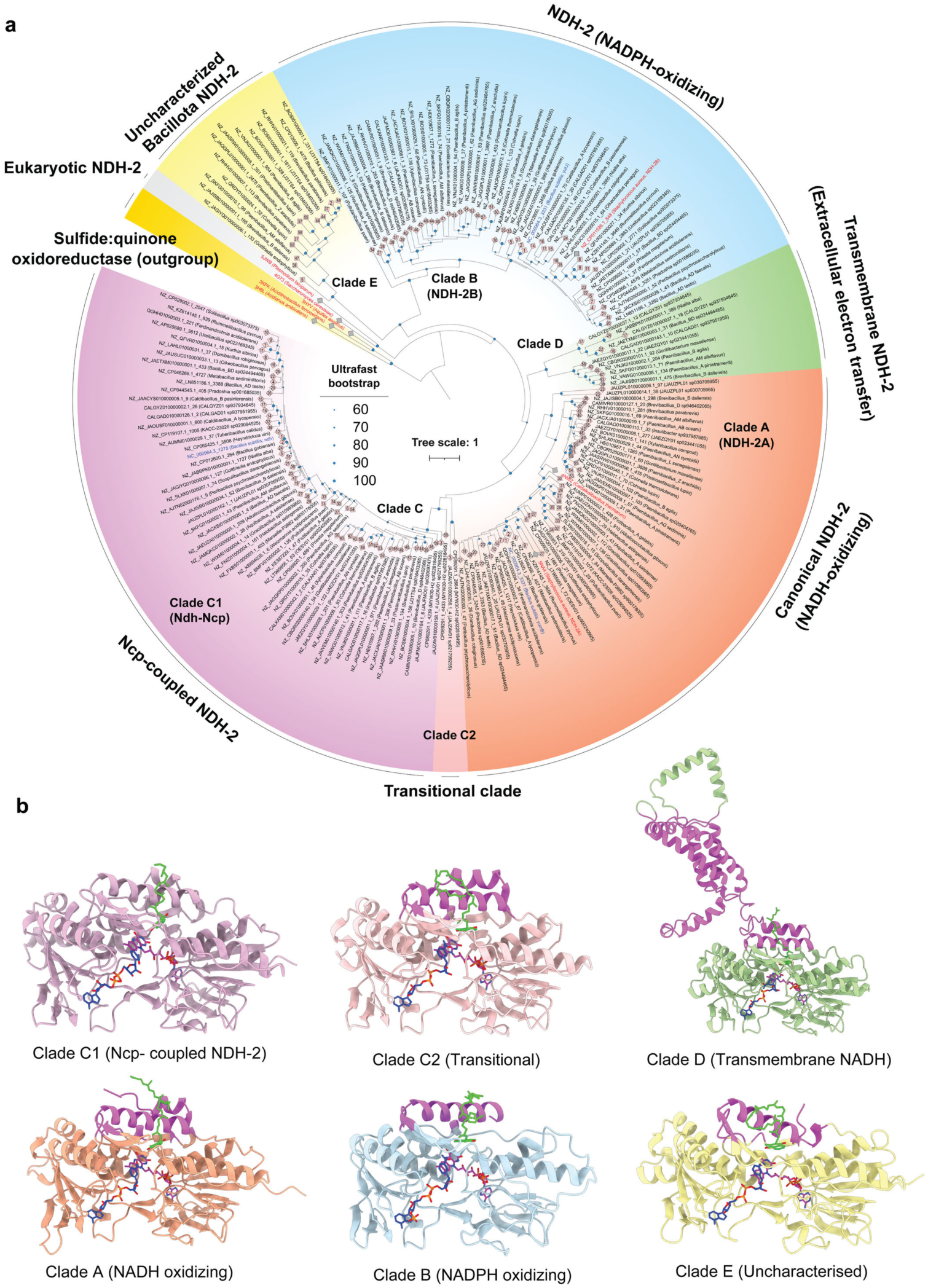
Ndh-Ncp is a novel and recently evolved NDH-2 clade widespread amongst Bacillota. a, A maximum-likelihood phylogenetic tree of NDH-2 from Ndh-Ncp encoding Bacillota genomes. Sequences are from a representative subset of 60 phylogenetically divergent Bacillota species (see **Figure S8a** and **Supplemental Data S2** for all Ndh-Ncp encoding species), highlighting the common presence of multiple divergent NDH-2 genes within the same organism, which likely mediate di_erent physiological functions. The tree was constructed based on 195 amino acid sequences using the LG+I+R10 model and rooted using sulfide:quinone oxidoreductases (Sqr) as an outgroup. Taxonomy of the genomes follows GTDB R09-RS220. Coloured tip labels show reference NDH-2/Sqr sequences (red) and the three *Bacillus subtilis* NDH-2 (blue). Branch circles show ultrafast bootstrap support values with 1000 replicates. Diamond labels at the tip denote identifiers of the 60 genomes. The scale bar corresponds to the expected number of substitutions per site. **b,** Chai1 predicted structural models for representatives of each NDH-2 clade are shown below and coloured to match the tree. Membrane interaction regions are coloured in purple. Clade C1: *B. subtilis* Ndh (NC_000964.3_1275) (experimental data), Clade C2: *MYW30-H2 sp022818495* (CP089291.1_2664), Clade D*: B. daliensis* (NZ_JAJISB010000001.1_475), Clade A: *B. subtilis* YumB (NC_000964.3_3321), Clade B: *B. subtilis* YutJ (NC_000964.3_3331) and Clade E: *B. daliensis* (NZ_JAJISB010000001.1_165). Model validation in **Figure S10**.

For comparison, clades A and B represent widespread NDH-2 subtypes that frequently co-occur and genomically colocalise (separated by 4–20 genes) across divergent Bacillota species (**Figure 5a,b, Figure S8a, S7b, Supplemental Data S2**). Clade A represents the canonical monotopic NADH-oxidising NDH-2, containing *yumB* of *B. subtilis* and the structurally resolved NDH-2 enzymes from *S. aureus* and *C. thermarum* (**Figure S9a,b,c**). These structures feature C-terminal helices that anchor NDH-2 at the membrane surface, facilitating quinone access (**Figure S9a,b,c**)^40–42,49^. Clade B is represented by the biochemically characterised monotopic NADPH-oxidising NDH-2 from *S. aureus* (NDH-2B)^49^, in addition to *yutJ* of *B. subtilis* (**Figure 5a,b, Figure S8a, Figure S9d**). Their genomic co-localisation suggests ancestral relationships and complementary functions in NADH and NADPH oxidation. Interestingly, clade D represents NDH-2 enzymes that associate with the membrane through transmembrane helices and has been implicated in extracellular electron transport (**Figure S9f**)^50,51^.

Because genomes containing *ndh–ncp* also typically encode clade A and clade B NDH-2 enzymes, we next asked why multiple NDH-2 subtypes coexist. Transcriptomic data from *B. subtilis* show that *ndh* (Ndh–Ncp) is expressed primarily during exponential growth, when respiratory activity is highest (**Figure S7c**)^52^. By contrast, *yumB* (clade A) is expressed during stationary phase and sporulation^35,52^, when energy demand is reduced, and fewer respiratory complexes occupy the membrane. *yutJ* (clade B) shows lower expression throughout, consistent with a specialized role in NADPH oxidation (**Figure S7c**)^35,52^. We hypothesize that the Ndh–Ncp complex functions as a space-efficient adaptation suited to exponential-phase respiration, when the membrane is crowded with respiratory complexes. In contrast, YumB employs surface-embedded helices, a mechanism compatible with the lower respiratory load of stationary phase (**Figure S9c**). These complementary modes of expression and membrane interaction underscore the distinct physiological roles of Ndh–Ncp and clade A NDH-2, a division of labour that appears conserved across Bacillota.

## Discussion

Our findings show that *B. subtilis* Ndh, the primary NADH dehydrogenase in this organism, forms a stable complex with the DUF1641 protein Ncp, which enables filament formation and quinone-mediated electron transfer to the respiratory chain (**Figure 6**). A defining feature of this complex is its large internal chamber, which accommodates numerous phospholipids, creating a solvent-excluded hydrophobic lumen that mimics the membrane bilayer. We observe Ndh–Ncp filaments of up to thirteen units, with chambers connected into a continuous hydrophobic conduit. This arrangement allows 52 Ndh subunits to access quinones with a membrane footprint similar to a single NDH-2 dimer embedded in the bilayer. These pseudomembrane filaments extend NADH-driven quinone reduction into the cytosol, expanding metabolic capacity by increasing the surface area available for respiration. This structure provides an elegant solution to the constraint of limited membrane surface area^1,2^. In addition, the ability of Ndh-Ncp to sequester quinone may also enable these filaments to function as a “quinone battery,” concentrating quinone for reduction by Ndh, which can be exchanged during transient interaction with the membrane. These features distinguish Ndh–Ncp from other DUF1641-containing enzyme complexes like Huc and ForCE, which incorporate few or no observable phospholipids and do not form filaments^23,24^.

**Figure 6.**
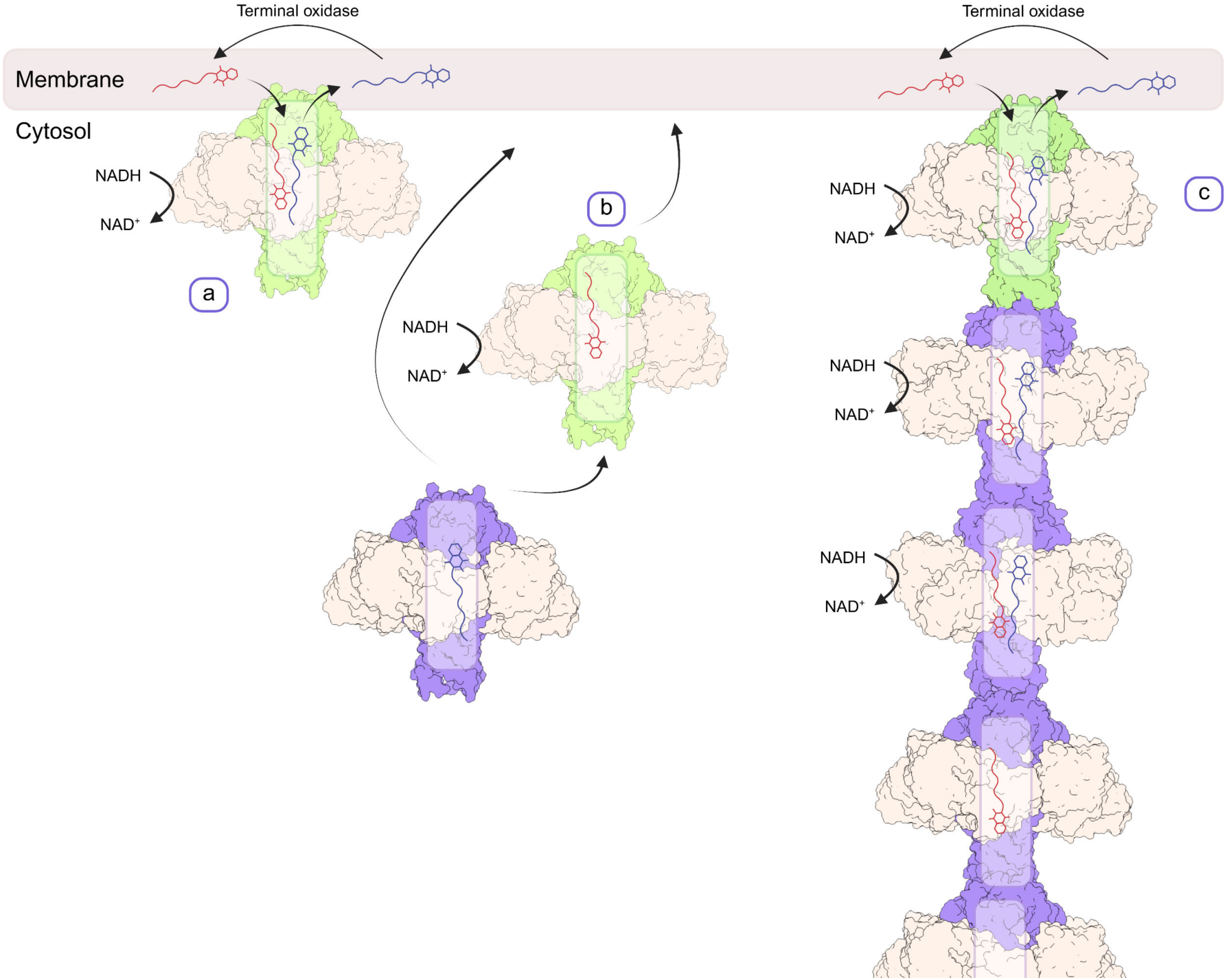
A model for quinone transfer and reduction by the Ndh-Ncp complex. a, Ndh can interact with the respiratory membrane through the C-terminal head region of the Ncp. The Ncp sequesters MK-7 (menaquinone in red) from the membrane for delivery to the electron acceptor sites in Ndh, where MK-7 is reduced (menaquinol in blue) by electrons from NADH oxidation. **b,** Ndh-Ncp can either remain at the membrane or dissociate using the MK-7 stored in the tube. Ndh-Ncp associates with other Ndh-Ncp complexes or directly with the membrane using the same interface. **c,** Ndh-Ncp forms a filament allowing for MK-7 to di_use from the membrane through several Ncp tubes to the associated Ndh-Ncp chambers for storage and reduction. Figure made with BioRender.

These properties place Ndh–Ncp alongside other strategies to expand bioenergetic capacity. In chloroplasts and cyanobacteria, thylakoids increase membrane area for photosynthetic electron transfer; in mitochondria, cristae enlarge the inner membrane and concentrate oxidative phosphorylation machinery; and in nitrifying and anammox bacteria, intracytoplasmic membranes and organelles support nitrite and ammonia oxidation respectively^3,4,8–12^. Ndh–Ncp represents a conceptually distinct solution to this problem, a protein–lipid filament that substitutes for additional membrane surface area, by creating an internal hydrophobic tunnel for quinone transfer. This mechanism is structurally similar to electron-transferring redox enzyme filaments like hydrogen-dependent CO_2_ reductase (HDCR), yet functionally more analogous to classical membrane elaborations^53^.

Physiological evidence supports the role of Ndh-Ncp in expanding respiratory capacity. Ndh-Ncp is the predominant NADH dehydrogenase during exponential growth, when respiratory demand is highest and the membrane is crowded by proteins supporting growth-related cellular processes^52^. The distribution of Ndh–Ncp across diverse Bacillota further supports the utility of this complex, though recent in evolutionary terms, it has been selected as an efficient strategy to enhance respiratory flexibility. Beyond its biological role, the Ndh–Ncp complex provides new design principles for synthetic bioenergetic systems. The ability to sequester and transport hydrophobic electron carriers using soluble protein-based filaments expands possibilities for building synthetic respiratory systems with greater spatial control of electron transfer than classical bilayers.

In summary, the Ndh–Ncp complex expands the conceptual and mechanistic framework of respiratory biology. By demonstrating that respiratory enzymes can operate through protein-based pseudomembrane filaments, this work reveals a mechanism for spatially separating respiratory redox chemistry from the membrane. This strategy alleviates the challenge of limited membrane surface area while preserving efficient quinone reduction. Future work should determine the quinone capacity of these complexes, how their filaments behave in vivo, and how they integrate with other respiratory systems. Together, our results establish Ndh–Ncp as a striking example of how evolution has reimagined membrane function and provide a foundation for engineering synthetic energy-transducing systems.

## Materials and methods

### Data analysis and visualization

In addition to programs described elsewhere in the manuscript, Microsoft Excel 2016, GraphPad Prism 10 and Origin were used for data analysis and visualization.

### Cloning and mutant construction

The *ndh* and *ncp* genes were amplified from *B. subtilis* 168 genomic DNA (gDNA) as one continuous fragment and subcloned into a pET22b plasmid, between BamHI and NheI sites, modified to encode an N-terminal twin strep II tag, yielding the expression vector p22b-Ndh-Ncp.

To generate isogenic deletion mutants, gDNA was isolated from the *B. subtilis* kanamycin gene deletion library strains Δ*yjlC*::kan (now Δ*ncp*::kan) and Δ*ndh*::kan as previously described^54,55^. Subsequently, our *B. subtilis* 168ca (trp+) wild-type strain was transformed with the isolated gDNA and mutants were selected on kanamycin-containing nutrient agar plates^54^. Colonies were clean-streaked on fresh nutrient agar-kanamycin plates and stored at-80 °C. Deletions were confirmed by PCR and Sanger sequencing. All primers, strains and plasmids are listed in **Supplemental Table S2**.

### *B. subtilis* growth assay

*B. subtilis* 168ca (trp+) wild-type^56^ and isogenic mutants Δ*ncp*::kan and Δ*ndh*::kan were routinely grown overnight in lysogeny broth (LB, 10 g L^-1^ tryptone, 5 g L^-1^ yeast extract, 10 g L^-1^ sodium chloride) at 37°C with aeration, with the addition of kanamycin at a final concentration of 5 µg ml^-1^ for the deletion mutants. The following day, cultures were normalised to a starting OD_600_ 0.05 in fresh LB and transferred in 200 µl aliquots, in technical triplicate and biological duplicate, to a 96-well plate, at either 37°C or 30°C. Growth was subsequently monitored at OD_600_ every 15 minutes for 12 hours in a BMG Spectrostar Nano plate reader. Growth rates were calculated using values between 0.1 and 1.6 OD_600_, equating to the bacterial growth during exponential phase. The rate of growth was calculated for each replicate and plotted as three individual data points. Growth yield experiments were carried out in LB + 0.5% glucose to reduce sporulation, and was calculated as OD_600_ reading at 24 hours after inoculation.

### Expression of Ndh-Ncp

p22b-Ndh-Ncp was transformed into *E. coli* C41 cells and grown as a starter culture in LB media until the culture reached the stationary phase, around 12-16 hours. Large cultures of 500 ml terrific broth media (TB, 12 g L^-1^ tryptone, 24 g L^-1^ yeast extract, 12.26 g L^-1^ K_2_HPO_4_, 2.31 g L^-1^ KH_2_PO_4_, 4 g L^-1^ glycerol) in a 2.5 L flask were inoculated with 5 ml of the starter culture and grown shaking, at 180 RPM, at 37°C until they reached ∼1.0 OD_600_. The cultures were then cooled at 4°C for 30 minutes before being returned to an incubator at 20°C and induced with 0.3 mM IPTG. The cultures were incubated for a further 16-18 hours before harvesting. Cells were harvested by centrifugation at 3500*g* for 20 minutes at 4°C using a Sorvall high-speed centrifuge. Once pelleted, cells were frozen and stored at-80°C.

### Purification of Ndh-Ncp

Purification occurred in a two-step process, beginning with affinity purification and followed by size exclusion chromatography (SEC). Initially, the cells were thawed and resuspended with lysis buffer (50 mM Tris, 150 mM NaCl, pH 8.0, 1× cOmplete EDTA-free protease inhibitor (Roche, 11836145001), 1 mg ml^−1^ DNase and lysozyme) at room temperature. Cells were then lysed by two passages through a cell disruptor (Emulsiflex C-5), and the lysate was clarified by centrifugation at 30,000*g* for 20 minutes at 4°C. Lysate was then incubated for 5 minutes with BioLock biotin blocking buffer (IBA Lifesciences IBA, 2-0205-050) and loaded onto a 1 ml StrepTrap XT (Cytiva). Once loaded, the column was washed with lysis buffer for ∼ 30 column volumes (CV). The protein was eluted with elution buffer (50 mM Tris, 150 mM NaCl and 50 mM biotin, pH 8.0) in ten 1 ml fractions. Resulting fractions were analysed by SDS-PAGE, then pooled and concentrated to 500 µl for size exclusion chromatography (SEC) using a 4 ml centrifugal concentrator with a 100-kDa molecular weight cutoff (MWCO) (Amicon, UFC803008). Concentrated protein was then loaded onto a Superose 6 Increase 10/300 GL column (Cytiva, 17517201) in SEC buffer (50 mM Tris, 150 mM NaCl, pH 8.0) and eluted in 0.5 ml fractions. Fractions were analysed by SDS-PAGE for peak identification and purity. Peaks corresponding to the correct complex were either pooled and concentrated for biochemical assays or individual fractions were concentrated to separate each region of the peak for cryo-EM.

### Visible-wavelength spectroscopic measurements

These measurements used a (SpectraMax ABS Plus, Molecular Devices) 96-well plate reader fully housed in an anaerobic glovebox (O_2_ < 3 ppm; Belle Technology Ltd, UK). The enzyme was carefully deoxygenated, diluted with anaerobic buffer (50 mM Tris-Cl pH 8, 150 mM NaCl), and spectra were recorded in a quartz cuvette following the addition of NADH (0-100 µM). Data were plotted using GraphPad Prism.

### Steady-state oxidation of NADH measurements

The oxidation of NADH was measured at 340-380 nm (ε = 4.81 mM^−1^ cm^−1^) in a (SpectraMax ABS Plus, Molecular Devices) 96-well plate reader. The buffer (50 mM Tris-Cl pH 8, 150 mM NaCl) contained 0.15% asolectin-CHAPS to provide a hydrophobic phase for the short-chain quinones. Menadione reaction kinetics were measured with the addition 4 nM Ndh-Ncp at a concentration range of 0-120 µM menadione or 0-500 µM NADH. Decylubiquinone was measured with the addition of 8 nM Ndh-Ncp, over a concentration range of 0-120 µM decylubiquinone or 0-500 µM NADH. The data were fit to the Michaelis-Menten equation (*v* = V_max_ × [S] / (*K*_M_ + [S]) using the Solver in Microsoft Excel. Data were plotted in GraphPad Prism for figures.

### Negative stain and Cryo-EM imaging and data collection of Ndh-Ncp

Negative stain grids (carbon film 300 mesh, EMS) were prepared by first glow discharge for 30 seconds at 15 mA, and then the protein was applied (0.005 mg ml^-1^) for 1 minute, followed by blotting of the excess protein. Three rounds of uranyl acetate application followed (two at 30 seconds, the final for 1 minute, each followed by blotting). Grids were then imaged in a Talos L120C (Thermo Fisher).

Cryo-EM grids were prepared using a Vitrobot Mark IV (ThermoFisher Scientific) pre-cooled to 4°C and equilibrated to 100% humidity. UltrAuFoil 1.2/1.3 300 mesh grids (Quantifoil) were glow-discharged for 180 seconds at 15 mA. 4 µl of protein was applied at 6 mg ml^-1^ concentration to the grid after glow discharge. The grid was then plunge frozen into liquid ethane using the Vitrobot. Grids were immediately clipped for the autoloader and stored in liquid nitrogen long-term.

Samples were initially screened on a Talos Arctica 200 kV microscope, collecting a small dataset (200-300 movies) to generate 2D class averages with cryoSPARC live^57^. Large datasets were collected on a Titan Krios G4 fitted with a Gatan K3 direct electron detector (Ndh-Ncp) or Falcon4i direct electron detector (Ndh-Ncp NADH/quinone bound). Data were collected at 130,000 magnification (0.655 pixel size) with a total dose of 39 e^-^ per Å^2^, fractionated into 40 frames, for Ndh-Ncp, and 96,000 magnification (0.808 pixel size) with an exposure time of 3 seconds and a total dose of 50 e^-^ per Å^2^, which was fractionated into 50 frames, for Ndh-Ncp NADH/quinone bound. A C2 aperture of 42 µM was used with the objective aperture inserted and a defocus range between-1.2 µM and-0.6 µM. Data collections were automated using the EPU software (Thermo Fisher Scientific).

### Cryo-EM data processing and analysis

#### i) Ndh-Ncp

7876 micrographs were grouped into their respective optics groups using the python script AFIS.py and then the grouped micrographs were motion corrected using UCSF MotionCor3 and CTF estimation (CTFFIND-4.1) in Relion 5.0^58^. The particle coordinates were then determined with crYOLO (1.7.6) using a general model and extracted in Relion with a box size of 512 px^58,59^. Initial 2D classification was carried out in Relion 5.0 then extracted and imported into cryoSPARC for ab initio reconstruction, with one class with C4 symmetry^57^. Homogeneous refinement of 483,556 particles was followed by non-uniform refinement with C4 symmetry. 3D classification followed with 6 classes. Four classes were selected containing 389,955 particles and subjected to another round of non-uniform refinement with C4 symmetry. This generated a map of 2.14 Å global resolution which was improved to 2.12 Å resolution with local refinement, masking only one unit of the filament. From this, helical refinement was carried out with a helical rise of 99.22 Å, twist of 23.95° and final resolution of 3.09 Å. Alternatively, the particles from the local refinement were reclassified with heterogeneous refinement, masking one class with ‘bad particles’. The resulting ‘good particles’ were reclassified in 2D space, and 356,549 particles were selected for local refinement, resulting in maps reaching a global resolution of 2.11 Å (**Figure S2**).

#### ii) Ndh-Ncp NADH/quinone bound

11,418 micrographs were grouped into their respective optics groups using the python script AFIS.py, and then the grouped micrographs were motion corrected using UCSF MotionCor3 in Relion 5.0. The motion corrected particles.star files were transferred to cryoSPARC v4.6.0 for contrast transfer function (CTF) estimation^57,58^. Particle coordinates were determined initially using Blob Picker to generate 2D averages to train Topaz Picking^60^. Topaz Picking was trained only on the 2D class of the top view, in an effort to find more of this underrepresented view. Topaz Picking resulted in 5,095,952 particles, which were extracted at a 420-pixel box size, binned to 120 px, and 2D classification resulted in 2,590,595 particles selected. Ab initio was carried out with two classes and C4 symmetry, with the largest class containing 1,301,383 particles used for homogenous refinement (C4). Homogeneous refinement was followed by heterogeneous refinement, with two classes, one masked with the ‘junk’ ab initio class, the other masked from homogenous refinement. 797,409 particles were used for non-uniform refinement to generate a new model for Topaz picking. 1,894,345 particles resulted and were extracted at a 420-pixel box size, binned to 120 px. 1,267,220 particles were selected from 2D classification and re-extracted without binning. 2D classification further resulted in 1,217,300 particles selected and ab initio was carried out with two classes and C4 symmetry. Homogenous refinement (C4) with 632,082 particles resulted in maps to a global resolution of 2.58 Å. Further non-uniform refinement and 3D classification using heterogeneous refinement (two rounds) resulted in a non-uniform refinement with 341,297 particles to a global resolution of 2.52 Å (**Figure S3**).

### Model building and visualisation

An initial model of the Ndh-Ncp complex was generated with AlphaFold3^61^, and Ncp and Ndh were then separated into individual molecules using Pymol^62^. The Ncp tetramer and Ndh enzymes were docked separately into the maps generated, using ChimeraX fit map^63^. The models were built and refined using Coot and Phenix^64,65^. FAD, MK-8 and NADH were modelled into the complexes using Coot^65^. Model quality was validated using MolProbity^66^. All figures were made using ChimeraX^63^.

### Mass spectrometric lipid detection

Ndh-Ncp purified in *E. coli* was incubated with *B. subtilis* 168 membranes extracted from 4 L of *B. subtilis* cultures or left untreated. To prepare the membranes, *B. subtilis* was grown in LB at 30°C for 24 hours, then harvested by centrifugation at 3500*g* and frozen at-20°C. Cells were lysed in buffer (50 mM Tris, 150 mM NaCl, 0.5 mM EDTA, pH 7.5) and incubated with 1 mg ml^-1^ lysozyme at 37°C for 30 minutes. Cells were then incubated with 1× cOmplete EDTA-free protease inhibitor (Roche, 11836145001) and 1 mg ml^−1^ DNase for ten minutes at room temperature, followed by lysis with two passages through a cell disrupter (Emulsiflex C-5). The lysate was clarified by centrifugation at 27,000*g* for 20 minutes at 4°C, and then the membranes were harvested by ultracentrifugation at 100,000*g* for 1 hour at 4°C. Membranes were resuspended in 50 mM Tris, 150 mM NaCl, pH 7.5 and incubated for 16 hours at 4°C with 5 mg of Ndh-Ncp rotating slowly. The membranes were then pelleted by ultracentrifugation, and the resulting protein was purified by SEC on a Superose 6 Increase 10/300 GL column (Cytiva, 17517201) in SEC buffer (50 mM Tris, 150 mM NaCl, pH 8.0). Resulting eluted fractions were pooled, concentrated and frozen for mass spectrometry.

Samples corresponding to 227.5 µg of purified protein (determined by absorbance at 280 nM) were prepared for liquid chromatography-mass spectrometry (LC-MS) analysis using a modified Folch extraction. Briefly, a 50 μl solution of purified protein (∼227.5 μg) was treated with 1000 μl of 2:1 chloroform:methanol v/v after which the mixture was shaken for 10 mins and allowed to stand for a further 50 mins. 200 μl of water was added, and the mixture was shaken for 10 minutes after which the sample was allowed to stand until the two phases had completely separated. The lower chloroform-rich phase was then transferred to a 1.4 ml total recovery glass microvial (ThermoFisher Scientific PN 6PSV9-CHV1) 2 ml sample vial, and the solvent was removed under a stream of nitrogen gas. Samples were analysed using a Vanquish Horizon (Thermo Fischer Scientific, Australia) coupled to a Q-Exactive Plus Orbitrap MS (Thermo Fischer Scientific, Australia) using a C18 column (Zorbax Eclipse Plus C18 Rapid Resolution HD 2.1 x 100mm 1.8 micron, Agilent, 959758-902) with a binary solvent system; solvent A = 40% isopropanol and solvent B = 98% isopropanol both containing 2 mM formic acid and 8 mM ammonium formate. Linear gradient time-%B as follows: 0 min-0%, 8 min-35%, 16 min-50%, 19 min-80%, 23 min-100%, 28 min-100%, 30 min-0%, 32 min-0%. The flow rate was 250 μl min^-1^, the column temperature 50°C, and the sample injection volume was 10 μl. Each sample was analysed three times, firstly by at an MS1 level using fast polarity switching. Then by data-dependent acquisition (DDA) in a single polarity, ie. one analysis in positive mode and one analysis in negative mode. For the polarity switching analysis, the mass spectrometer operated at a resolution of 140,000 from m/z = 140 to 2000 and DDA was performed with a MS1 survey scan m/z = 450 to 2000 at a resolution of 70,000. The resulting data-dependent MS/MS was conducted at a resolution of 17,500 using an isolation window of 1.5 m/z and a stepped normalized collision energy 20, 40 and 60. The method cycled with a loop count of 7 and dynamic exclusion of 8.0 seconds. The following source conditions were used in all experiments: Spray voltage 4.0 kV (positive) 3.5 kV (negative); capillary temperature 300 °C; sheath gas 50; Aux gas 20; sweep gas 2 and probe temp 120 °C.

Lipidomic data analysis was performed using untargeted feature extraction on the MS1 polarity switching data in mzMine 4.3^67^. Briefly, this involved untargeted feature extraction from each sample, alignment of features and gap-filling and identification against those features previously identified. Features that did not exceed the signal found in the extraction blanks by at least 5-fold were excluded. Lipids were then putatively identified by comparison against the Lipid identification, which was achieved by comparing the MS/MS spectra against the LipidBlast database^68^ MSDIAL-TandemMassSpectralAtlas-VS69^69^m followed by manual review of the putative annotations. Unassigned features were then annotated based on their accurate mass against the Lipid Maps structural database^70,71^ (MS1ID) and or the MycoMass database^72^. Raw and processed data can be found in the accompanying repository (https://store.erc.monash.edu.au/experiment/view/20090/?token=AJK8L11I2TOAXNYKYEGTYXM6BBIOME)

*E. coli* purified Ndh-Ncp were analysed for abundance of lipids by adding the peak area of each category of lipid (PE, PG, UQ-8, MK-8, DMK-8) and calculating the relative peak area in the total sample. This was then normalised so the most abundant lipid (PE) was designated 100. For the *B. subtilis* incubation experiment, the peak area of representative lipids present in the sample before and then after extraction were calculated and then the fold change was calculated based on this (amount after incubation/before incubation).

### Estimation of quinone-to-phospholipid ratio in *E. coli* membranes

To estimate the bulk quinone:phospholipid ratio in the *E. coli* inner membrane, we compared reported absolute abundances of quinones and phospholipids. Biochemical analyses indicate that aerobic *E. coli* contains ∼0.4–1 µmol of respiratory quinone per g dry weight (DW)^73^. In contrast, lipids account for ∼9–10% of cellular dry mass, of which >90% are phospholipids^74,75^. Assuming an average molecular weight of ∼750 g·mol⁻¹ for phospholipids, this corresponds to ∼100–120 µmol phospholipid per g DW. On this basis, the expected quinone:phospholipid ratio is ∼1:100–1:300 under the aerobic conditions, where Ndh-Ncp was produced.

### Molecular dynamics simulations

We performed all-atom classical molecular dynamics (MD) simulations on the high-resolution cryo-EM structures of *B. subtilis* Ndh-Ncp complex with and without MK-7/NADH. Simulations were performed with GROMACS 2022.3/4^76^ using the CHARMM36m force field for proteins and CHARMM36 for lipids, ions, water, and redox-active components (FAD, MK-7, NADH)^77–80^. The membrane bilayer was constructed with the CHARMM-GUI^81^ tool, and comprised 950 POPE (∼79%) and 259 POPG (∼21%) lipids. The total system size was ∼664,000–666,000 atoms depending on lipid and substrate arrangements (see more below). In total, seven simulation setups were constructed (**Supplemental Table S3**).

Two strategies were applied for lipid placement in the Ndh-Ncp complex. In the first (setup 1), lipids were manually embedded into Ndh-Ncp interface cavities with VMD’s graphical interface^82^. The lipid head groups were oriented towards the solvent and tails directed inward, resulting in 33 POPE and 7 POPG molecules in a tightly packed chamber. In the second approach (setup 2), a pre-equilibrated bilayer patch was aligned using the *psfgen* plugin^82^ with Ncp and only lipids inside the chamber were retained, resulting in a total of 20 POPE and 7 POPG molecules (setups 2, 3, 4, 6, **Supplemental Table S3**). Lipids near the N-terminal region were added similarly.

All membrane-protein model systems were solvated with TIP3P water and 150 mM Na^+^/Cl^-^, using VMD’s *autoionize* and *solvate* plugins^82–84^. Protonation states were predicted using PROPKA at pH 7.0^85^. In all Ndh monomers, His383 was modelled protonated in non-substrate-bound setups, whereas Glu168 was protonated only in the MK-7/NADH-bound setups. Simulations were performed in the isothermal-isobaric (NPT) ensemble (310 K, 1 atm) with semi-isotropic pressure coupling. MD equilibration used the V-rescale thermostat and C-rescale barostat, followed by production simulations with Nosé–Hoover thermostat and Parrinello–Rahman barostat^86–90^. A 2 fs timestep was used with the leapfrog integrator, and all covalent bonds involving hydrogen atoms were constrained using the LINCS algorithm^91,92^. Nonbonded interactions were applied with a 12 Å cutoff and particle-mesh Ewald (PME)^93,94^ for long-range electrostatics.

The membrane patch was pre-equilibrated for 10 ns by applying harmonic position restraints (20,000 kJ mol^-1^ nm^-2^) to the phosphate groups of all lipids, followed by a 30 ns production run to generate a fully equilibrated lipid bilayer. Simulations of membrane-protein systems (**Supplemental Table S3**) began with a 1000-step energy minimisation using NAMD 2.14^95,96^, with the harmonic position restraints (force constant of ≈ 41.4 kJ mol^-1^ nm^-2^ applied to all heavy atoms of the protein and its substrates. This was followed by a steepest-descent minimisation in GROMACS 2022.3 with an energy tolerance of 250 kJ mol^-1^ nm^-1^, using restraints (20,000 kJ mol^-1^ nm^-2^) on the same atoms. Subsequently, equilibrations were performed in the NPT ensemble with identical position restraints for 100 ns to relax the lipids within the Ncp tube, followed by a 100 ns equilibration by keeping the same harmonic restraints now only on the protein backbone and FAD atoms. Three production simulation replicas were run per setup, where replicas 2 and 3 were generated by extending the last equilibration step by 1 and 2 ns, respectively. Additionally, in substrate-bound systems, weaker restraints (2000 kJ mol^-1^ nm^-2^) were applied to NADH and MK-7 headgroups during equilibration setups. Production simulations were extended to 1 µs. It is noteworthy that MK-7 molecules (setup 5), after binding to FAD for ca. 300 ns, diffused from their structural locations. Therefore, in additional simulations (setup 6, **Supplemental Table S3**), harmonic distance restraints were applied between FAD (N5) and NADH (C5N), and between NADH (C5N) and the MK-7 (OR8) atoms, using the GROMACS pull code with a force constant of 5000 kJ mol^-1^ nm^-2 76^. Volume calculations of solvent accessible surface (SAS) were performed with the MSMS program^97^ combined with the Openstructure software package^98^.

### Synthesis and characterisation of S-cholesterol (for electrochemistry membrane bilayer)

#### i) Amide Coupling

2-[2-(2-Aminoethoxy)ethoxy]ethanol **1** (1.08 g, 1.00 ml, 7.22 mmol) was dissolved in anhydrous DCM (25 ml) and triethylamine (0.82 g, 1.12 ml, 8.04 mmol) was then added. Separately, cholesteryl chloroformate (3.62 g, 8.04 mmol) was dissolved in anhydrous dichloromethane (DCM) (50 ml). The solution of cholesteryl chloroformate was then added dropwise to the solution of 2-[2-(2-Aminoethoxy)ethoxy]ethanol, and the resultant solution was stirred for 48 hours at room temperature under N_2_. After this time, the reaction mixture was diluted with additional DCM (100 ml) and was washed with 1 M HCl (2 x 40 ml), then brine (40 ml). The organic layer was dried over MgSO_4_ and was concentrated *in vacuo* to yield a crude residue, which was purified via silica column chromatography (hexane → ethyl acetate) to yield **2** as a colourless oily foam (3.95 g, 97%). Note, when performing TLC, that the location of **2** can be visualised with permanganate stain (**Figure S11a**).

#### ii) Preparation of Thioester

**2** (1.95 g, 3.47 mmol) was dissolved in anhydrous DCM (50 ml). Triethylamine (630 ul, 4.51 mmol) was then added, followed by p-Toluenesulfonyl chloride (0.860 g, 4.51 mmol). The reaction solution was then stirred at room temperature under N_2_ overnight. After this time, the reaction solution was diluted with additional DCM (50 ml) and washed with 1 M HCl (2 x 20 ml), sat NaHCO_3_ (20 ml) and then brine (20 ml). The organic layer was dried over MgSO_4_ and concentrated *in vacuo* to yield the crude tosylate intermediate. The resulting residue was then re-dissolved in anhydrous dimethylformamide (DMF) (20 ml) and potassium thioacetate (1.19 g, 10.4 mmol) was then added. The resultant solution was then placed under a nitrogen atmosphere and was stirred at 90°C for 3 h, during which time a strong brown colour developed. The reaction solution was then cooled to room temperature and the majority of the DMF was removed *in vacuo*. DCM was then added (100 ml), and the resultant solution was transferred to a separating funnel. The solution was washed with water (2 x 50 ml) and then brine (20 ml) and was then dried over MgSO_4_. The product, **3**, was then isolated via silica column chromatography (10% EtOAc in petroleum ether), yielding a deep-brown oil (0.910 g, 42%) (**Figure S11b**).

#### iii) Deprotection of Thiol

**3** (0.882 g, 1.42 mmol) was dissolved in ethanol (10 ml). Separately, potassium hydroxide (169 mg, 3.01 mmol) was dissolved in ethanol (10 ml) and delivered to the solution of **3**. The reaction mixture was placed under a nitrogen atmosphere and was stirred at room temperature for 3 hours before being diluted in DCM (50 ml). The reaction mixture was then transferred to a separating funnel. The solution was washed with 1 M HCl (10 ml), then brine (20 ml), and was then dried over MgSO_4_. Concentration *in vacuo* yielded a crude brown residue, which was further purified via silica column chromatography (hexane → ethyl acetate). **4** was isolated in resolved fractions as both its free-thiol monomeric form (0.419 g, 51%) and its corresponding disulfide dimer (0.209 g, 26%), in a combined yield of 77% (both the free-thiol and disulfide forms of the product can be used identically for SAM formation). Both forms of **4** were brown in colour (**Figure S11c**).

### Electrode modification

Liposomes were formed from a solution of 5 mg ml^-1^ soy phospholipids (Sigma Aldrich) in aqueous buffer (20 mM MOPS, 30 mM Na_2_SO_4_, pH 7.40). This suspension was extruded through an Avanti mini-extruder containing a Cytiva 0.1 μm membrane filter thirty-one times to generate liposomes. When quinone supplementation was required, 0.44 mg ml^-1^ menaquinone-7 (“MK-7”, Cambridge Bioscience) or 0.44 mg ml^-1^ ubiquinone-8 (“UQ-8”, Merck) was added to the lipid suspension before extrusion.

BASi gold disc electrodes (3 mm diameter) were first mechanically polished in air using alumina slurries of decreasing particle size (1, 0.3, then 0.05 μm, Metprep) applied to polishing pads (Beuhler), with 2-minute polishes used per alumina size followed by sonication for 3 minutes in water, then ethanol. The electrodes were then taken into a glovebox (in-house construction, Belle Technology purification columns, [O_2_] < 30 ppm) and an electrochemical polish of 50 cyclic voltammograms from-1.3 to 0.7 V vs Ag/AgCl at 100 mV s^-1^ in 0.5 M NaOH were followed by 50 cyclic voltammograms from-0.25 to 1.55 V vs Ag/AgCl at 100 mV s^-1^ in 0.5 M H_2_SO_4_. Remaining in the glovebox, the cleaned electrodes were left to dry and then immersed in 1 ml of 2-propanol solution containing 8.9 mM 6-mercaptohexanol and 1.1 mM EO_3_-cholesterol (S-cholesterol) for 16 hours in order to form a mixed self-assembled monolayer (SAM), following the published procedure by Nakatani et al^47^. Following this SAM formation, the electrodes were removed from the glovebox and rinsed with ethanol and dichloromethane in the laboratory atmosphere and dried under N_2_. The electrodes were then returned to the glovebox and in order to form the liposome-functionalization, 5 μl liposome was pipetted onto the tip of the electrode, and 1 μl of 120 mM CaCl_2_ in the same buffer solution was added with gentle mixing using the pipette tip. This was left to dry (approx. 30 minutes), then the same liposome and CaCl_2_ dropcast plus drying process was repeated. Enzyme-modification of the electrode was then achieved by applying 5 μl of 7 mg ml^-1^ *Bacillus subtilis* Ndh-Ncp in 50 mM Tris, 150 mM NaCl, pH 7.4 to the electrode, and leaving this to dry for approximately 30 minutes.

### Electrochemical set up

All experiments were carried out in a 3-electrode cell containing 10 ml buffer solution (20 mM MOPS, 30 mM Na_2_SO_4_, pH 7.40), which was stirred continuously throughout. The cell used a modified gold disc working electrode (see above), a platinum wire counter electrode and a Ag/AgCl reference electrode, which was determined to be offset from SHE by +0.225 V based on calibration experiments using ferricyanide (**Figure S6f**)^99^. All experiments were carried out in a glove box under a nitrogen atmosphere ([O_2_] < 30 ppm) at 19°C. During an NADH titration experiment, the potential was continuously cycled at 10 mVs^-1^, with aliquots of a concentrated stock solution of 100 mM NADH added in increments at approximately-0.4 V vs SHE during every even scan. Every odd scan is displayed.

### Genomic survey, synteny characterization, and phylogenetic analysis

To profile the distribution and diversity of the Ndh-Ncp complex across the prokaryotic tree of life, we retrieved all 113,104 representative archaeal and bacterial species genomes from the Genome Taxonomy Database (GTDB) release 09-RS220^48^. Genes were predicted using Prodigal v2.6.3^100^ and functionally annotated against the KEGG^101^ (accessed 22 November 2021) and Pfam v34.0^102^ protein databases through DRAM v1.2.4^103^. DUF1641-domin family proteins (including Ncp) were identified by searching against the Conserved Domain Database (CDD)^104^ profiles pfam07849 (DUF1641-domain) and COG2427 (YjgD/ForE, DUF1641-domain protein) using rpsblast (-evalue 1e-5-max_hsps 1-max_target_seqs 5) in BLAST+ v2.16.0^105^, as well as against all entries of DUF1641 domain proteins from UniProt^106^ using DIAMOND v.2.1.10^107^. Hits were cross-compared and manually inspected to filter false positives. For further curation and the analysis of genetic context, five genes upstream and downstream of DUF1641-domain hits and their annotations by DRAM were retrieved. Type II NADH dehydrogenases (NDH-2) in the neighbourhood and across genomes were identified by matches to KEGG Orthology KO K03885. In total, 43976 and 5479 genomes were found to encode NDH-2 and DUF1641-domain proteins, respectively; 684 genomes from exclusively the phylum Bacillota encode the Ndh-Ncp complex wherein *ndh* and *ncp* are always genomically co-located next to each other (**Supplemental Data S2**).

Genomes and NDH-2 sequences of Bacillota were further analysed to investigate the potential physiological relevance and evolution of the Ndh-Ncp complex. The presence of major aerobic and anaerobic respiratory enzymes (including Complexes I-V and terminal oxidases) in Bacillota genomes was identified based on DRAM and KEGG annotation. A phylogenomic tree of the Ndh-Ncp encoding Bacillota was constructed by aligning the concatenated 120 GTDB bacterial marker genes using GTDB-Tk v2.4.0^108^, followed by a maximum likelihood phylogenetic reconstruction using IQ-TREE v2.4.0^109^ (model: WAG + G20; 1000 ultrafast bootstrap replicates; mid-point rooted). For phylogenetic analysis of Bacillota NDH-2, Ncp-associated and non-Ncp-associated sequences with lengths over 300 amino acids were respectively clustered at identity thresholds of 80% and 60% using MMseqs2^110^ (-c 0.7 --cov-mode 1 --cluster-mode 2-s 7.5). A multiple sequence alignment of NDH-2 cluster representatives, structurally resolved NDH-2 (PDB: 5NA4, 4NWZ, 4G73, 5JWB), and sulfide:quinone oxidoreductases (PDB: 3H8L, 3HYV, 3KPK) was built using the align function of Muscle v5.3^111^. Using ModelFinder^111,112^ implemented in IQ-TREE v2.4.0, the best-fit model according to Bayesian information criterion (BIC) was determined to be LG+I+R10. A maximum likelihood phylogenetic tree was then constructed using IQ-TREE v2.4.0 (-m LG+I+R10-B 1000-bnni) and rooted using sulfide:quinone oxidoreductases as outgroups (**Figure S9**). To visualize the common presence of multiple copies of NDH-2 of different clades within the same microorganism, a maximum likelihood phylogenetic tree using all NDH-2 sequences from a subset of 60 representative and phylogenetically divergent Ndh-Ncp encoding Bacillota species was also constructed with the same method (**Figure 5**; **Supplemental Data S2**). All phylogenetic trees were visualized using iTOL v7^113^. Major clades of NDH-2 were defined in this study based on phylogeny, structural features (e.g. transmembrane domain, interaction with Ncp), and reported physiological roles (e.g. NADPH/NADH preference, extracellular electron transfer). NDH-2 and Ncp sequences, synteny of Ndh-Ncp, NDH-2 classification, genome statistics and metabolic annotations of the Bacillota genomes were summarized in **Supplemental Data S2**.

### Structural modelling

Modelling was performed on the Chai1 or AlphaFold3 online server^61,114^. Entire sequences were provided along with NADH (or NADPH)/FAD/MK-7/MK-8/UQ-8 smiles string.

## Data Availability

The cryo-EM structures are available in the PDB and EMDB under the codes PDB: **9PXK**, **EMD-71972** (Ndh-Ncp with MK-8/NADH), PDB: **9PXL**, **EMD-71973** (Ndh-Ncp) and PDB: **9PXM**, **EMD-71974** (Ndh-Ncp filament). Raw and processed lipidomics data can be found in the accompanying repository (https://store.erc.monash.edu.au/experiment/view/20090/?token=AJK8L11I2TOAXNYKY <u>EGTYXM6BBIOME</u>).

## Author Contributions

R.G., A.K., and K.A. conceived and designed the project. R.G. supervised the project. A.K. and K.A. expressed and purified proteins. A.K., K.A., D.R.F. prepared cryo-EM grids and collected data. A.K., R.G., and KA processed cryo-EM data. A.K. and R.G. modelled and analysed the structures. R.L.D. generated *B. subtilis* mutants and performed growth assays. J.A.S. and A.P. designed, performed and analysed cyclic voltammetry experiments. N.D.J.Y. and J.A.S. generated S-cholesterol for bilayer construction. J.N.B. and B.G.H. devised, performed and analysed UV-Vis and kinetics experiments. A.K. designed and carried out lipid exchange experiments, and C.K.B. designed, performed and analysed mass spectrometry experiments. O.Z., L.S., and V.S. devised, performed and analysed molecular dynamics simulations. P.M.L performed genomic surveys and phylogenetic analysis. A.K. and R.G. wrote and edited the manuscript with input from all authors.

## Acknowledgements

We thank Dr. Sepideh Valimehr and Dr. Hamish Brown from the Ian Holmes Imaging Centre for Cryo-EM training and advice. We thank Dr. Sarah Revitt-Mills from the Lyras lab at Monash University for providing the *B. subtilis* strain for lipid exchange experiments. We acknowledge the use of instruments at the Ian Holmes Imaging Centre. A.K., R.G, and D.R.F are members of the Australian Research Council Industrial Transformation Training Centre for Cryo-Electron Microscopy of Membrane Proteins for Drug Discovery (IC200100052). This work was supported by an ARC Discovery project grant awarded to R.G. (DP260100290). A.K. and D.R.F were supported by an RTP scholarship. This work was supported by a UKRI Future Leader Fellowship to J.N.B (MR/Z000084/1 & MR/T040742/1), a BBSRC sLoLa award to H.S. and J.N.B (BB/X003035/1), an Australian Research Council DECRA fellowship to P.M.L. (DE250101210), and an EPSRC PhD studentship to B.G.H (2885393). Funding for J.A.S., N.Y., and A.P. was provided by UKRI, EP/X027724/1. This study used BPA-enabled (Bioplatforms Australia) / NCRIS-enabled (National Collaborative Research Infrastructure Strategy) infrastructure located at the Monash Proteomics and Metabolomics Platform. This work was further supported by Monash eResearch capabilities, including Research Data Storage, M3 Massive HPC, and Nectar Research Cloud. VS acknowledges research funding from the Research Council of Finland, the Sigrid Jusélius Foundation, the University of Helsinki, the Jane and Aatos Erkko Foundation and the Finnish Society of Sciences and Letters. OZ acknowledges research funding from the Research Council of Finland. LS acknowledges support from the CHEMS doctoral programme of the University of Helsinki. We acknowledge extensive HPC resources from the Center for Scientific Computing (CSC), Finland and thank the IT support of the Faculty of Science, University of Helsinki.

## Supplemental Data

Supplemental data S1. Lipid identification output of purified Ndh-Ncp and *B. subtilis*

**membrane treated vs. untreated Ndh-Ncp**

**Supplemental data S2. Genomic survey results of NDH-2 from Bacillota**

## Supplemental Movies

**Supplemental Movie S1. Cryo-EM structure of filamentous Ndh-Ncp**

**Supplemental Movie S2. Density found associated with the Ncp tube corresponding to lipids**

**Supplemental Movie S3. Cryo-EM structure of Ndh-Ncp with NADH/MK-7 bound**

## Supplemental Notes

**Supplemental note 1. Preparation of the Ndh-Ncp-phospholipid complex MD simulations.**

The Ncp–Ndh-phospholipid complex was investigated using classical atomistic molecular dynamics simulations on two cryo-EM structures of the Ndh–Ncp complex with and without substrate. Guided by cryo-EM and lipidomics data, we examined the architecture and dynamics of the lipidic interior of the tube. The FAD, NADH, and MK-8 molecules associated with the complex were restrained during these simulations to generate a mode of the holo-Ndh-Ncp-phospholipid complex (see methods).

Volume estimates suggested the Ndh-Ncp chamber could accommodate ∼40–45 lipids. Accordingly, we generated a model with 40 lipids tightly packed into the Ndh-Ncp chamber and analysed its dynamics (**Figure S5a,c, setup 1**). In this configuration, lipid movement was highly constrained; which is not fully consistent with the blurred cryo-EM density observed in this region, indicating lipid mobility. We therefore constructed an alternative model where we aligned the pre-equilibrated lipid bilayer patch with the centre of the chamber, and retained the lipids populating the tube interior, resulting in 27-lipids bilayer inside the tube (**Figure S5a, setup 2**). Lipids in this new setup 2 were substantially more mobile than in setup 1 (**Figure S5c**). During microsecond-scale dynamics of setup 2, lipid head groups and hydration shells migrated toward the protein surface, exposing them to solvent, while water was expelled from the conduit, leading to a hydrophobic environment in the interior of the Ndh-Ncp complex (**Figure S5d-f**). By contrast, lipids in setup 1 remained largely immobile. These results indicate that the poorly resolved cryo-EM density is best represented by the smaller number of lipids modelled in setup 2, which we used as the basis for all subsequent simulations.

Simulations also revealed an abrupt conformational change at the N-terminal end of the Ncp tube, where partial collapse closed the opening. This is unlikely to be physiological, especially considering this region is associated with the C-terminal end of a neighbouring Ncp upon filament formation. To prevent this, we introduced five lipids into the region, which stabilised it in a conformation closer to the cryo-EM structure (**Figure S5a, setup 3**). The addition of these lipids left unoccupied space between the N-terminal region of Ncp and the main chamber of the Ndh-Ncp complex, which we filled with five additional, all lipids oriented tail-first toward the tube end (**Figure S5a, setup 4**). During equilibration of this model, ∼1–2 lipids spontaneously flipped, reorienting their head groups toward solvent while expelling water from the chamber. These simulations resulted in a stable model of the Ndh-Ncp-phospholipid complex, containing 30 lipids in the chamber and 5 at the N-terminal end (**Figure S5a, setup 6**), which was used for a final round of 1 μs (x3) simulations resulting in the model discussed in this paper.

## Supplemental figures

**Supplemental Figure 1.**
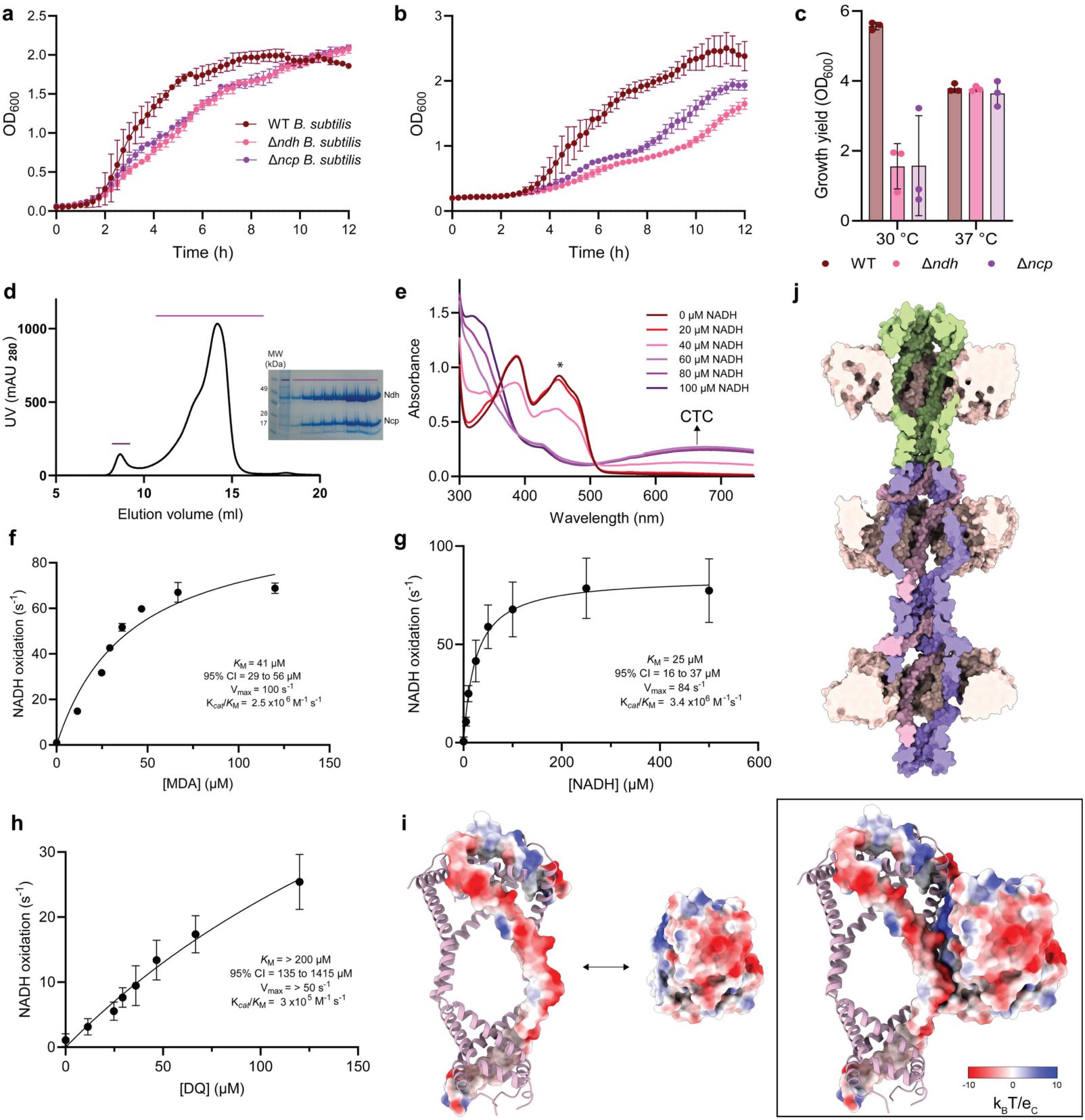
***B. subtilis* Ndh-Ncp purification and characterisation. a**, A growth curve of WT, Δ*ndh* and Δ*ncp B. subtilis* 168 strain, grown at 37°C, showing OD_600_ vs. time (h). **b,** A growth curve of WT, Δ*ndh* and Δ*ncp B. subtilis* 168 strain, grown at 30°C, showing OD_600_ vs. time (h). **c,** The growth yield of WT, Δ*ndh* and Δ*ncp B. subtilis* 168 strain after 24 hours of growth at 30 or 37 °C, measured in OD_600_. Data in panels a, b and c are from three biological replicates (n = 3) with mean ± standard deviation displayed. **d,** The SEC profile (Superose 6 10/300) from the purification of Ndh-Ncp with the corresponding SDS-PAGE on the right of the SEC curve. Ndh has an expected size of 42 kDa and Ncp 19.4 kDa (with strep tag). **e,** Representative UV-Vis spectra of Ndh-Ncp complex with increasing concentrations of NADH (all concentrations shown). **f,** Michaelis-Menton kinetics of Ndh-Ncp incubated with NADH and varying concentrations of MDA. **g,** Michaelis-Menton kinetics of Ndh-Ncp incubated with decylubiquinone (DQ) and varying concentrations of NADH. **h,** Michaelis-Menton kinetics of Ndh-Ncp incubated with NADH and varying concentrations of DQ. **i,** An electrostatic surface representation of the cryo-EM structure of one Ncp and one Ndh overlayed on the cartoon representation of NCP tube showing the interaction interface between one Ncp and one Ndh. Red colouring corresponds to negatively charged residues, white neutral, and blue positively charged residues. **j,** Surface representation of the Ndh-Ncp cryo-EM structure cut down the centre to show the continuous Ncp tube between complexes. Data in panels f, g and h are from three independent replicates (n = 3) with mean ± standard deviation displayed.

**Supplemental Figure 2.**
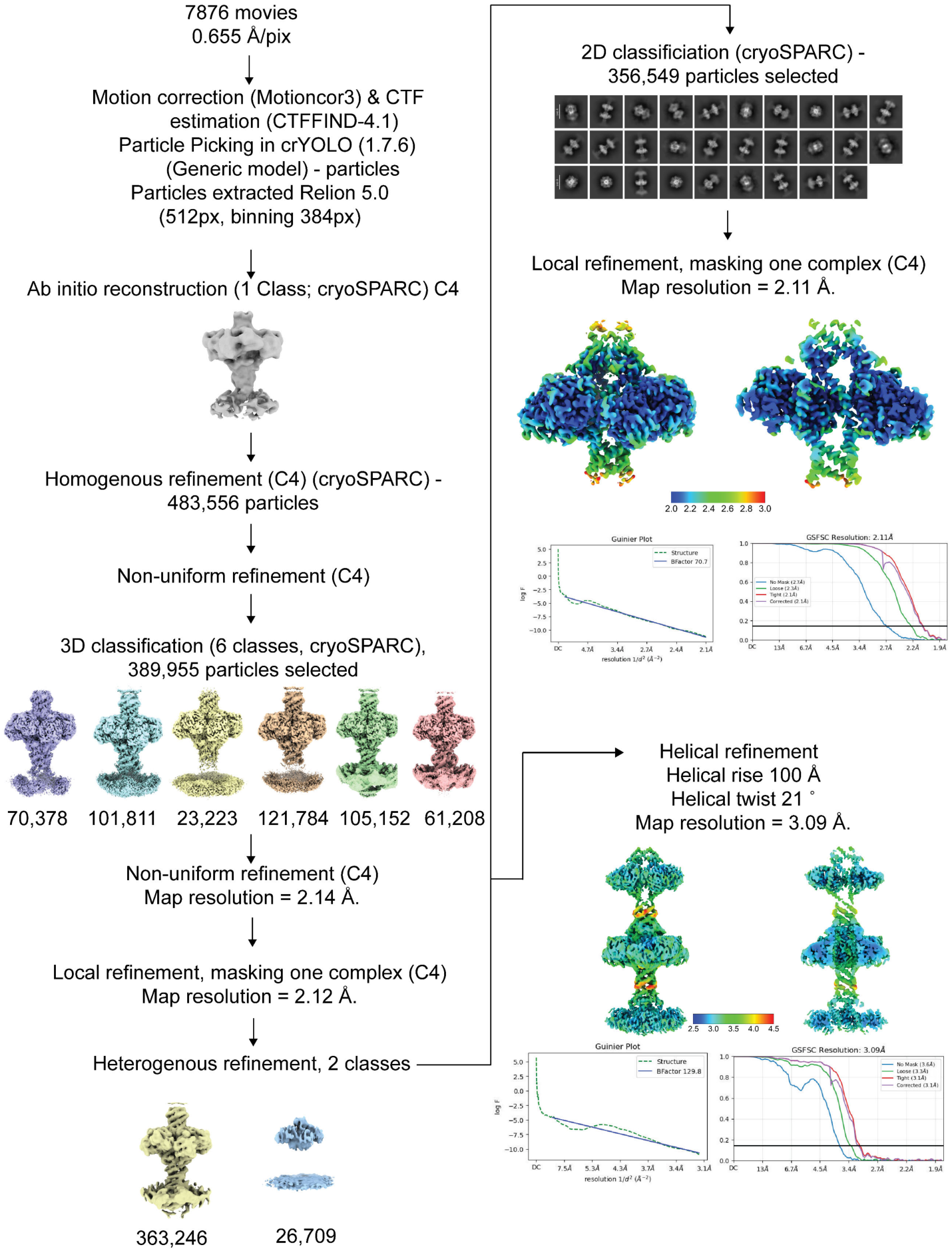
Ndh-Ncp cryo-EM data processing workflow. Beginning with 7876 movies, the data was processed in Relion 5.0, crYOLO and cryoSPARC, resulting in two maps, one for the filament at 3.09 Å global resolution and one Ndh-Ncp unit at 2.11 Å global resolution.

**Supplemental Figure 3.**
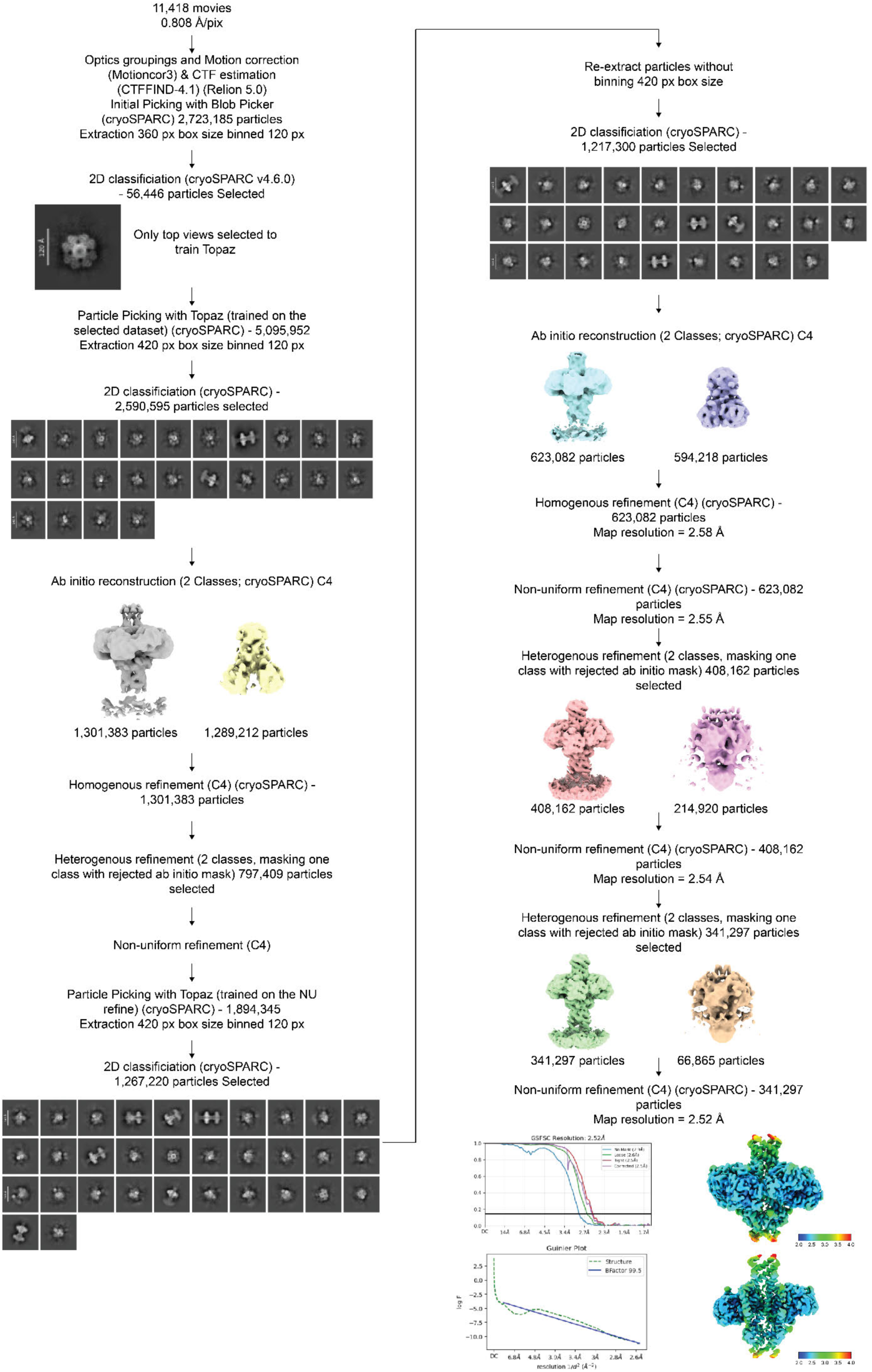
Ndh-Ncp NADH-MK-8 data processing workflow. Beginning with 11418 movies, the data was processed in Relion 5.0 and cryoSPARC, resulting in a map of 2.52 Å.

**Supplemental Figure 4.**
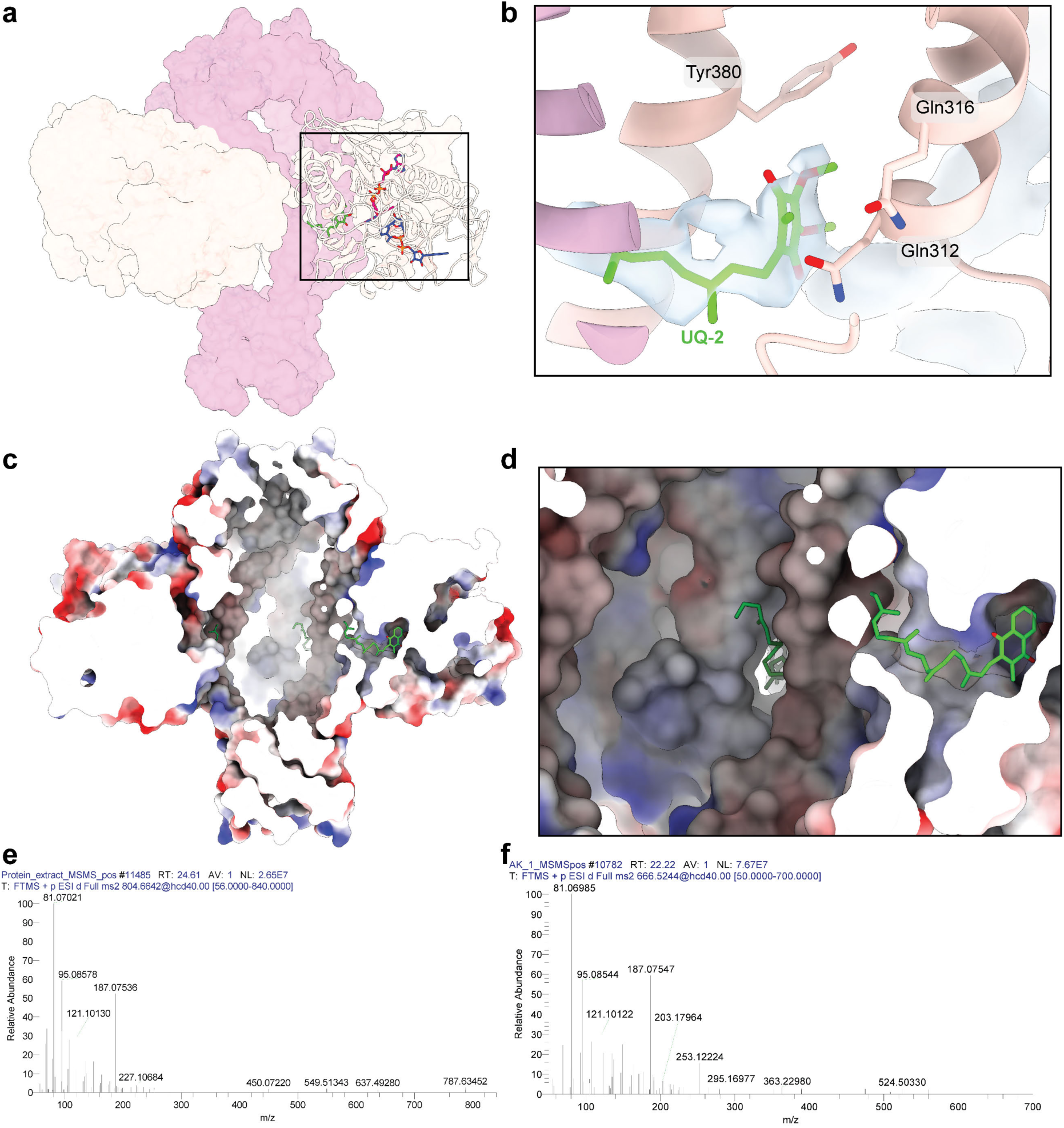
Ncp quinone and lipid association. a and b, Cartoon representation of Ndh-Ncp with UQ-2 modelled into the quinone binding site (green). Density is shown around UQ-2. **c,** Electrostatic surface representation of Ndh-Ncp cut away down the middle vertically showing the internal tube and the MK-8 molecules interfacing into the tube. **d,** A zoom of a MK-8 molecule (with the tail length truncated based on density) showing the entrance to the Ndh quinone binding site from the Ncp tube. **e and f,** Base peak chromatograms for the high-performance liquid chromatography–mass spectrometry analysis of Folch extracts from Ndh-Ncp after membrane incubation from positive ion mode. **e** Shows the MQ-9 standard, and **f** shows the molecule identified as MK-7.

**Supplemental Figure 5.**
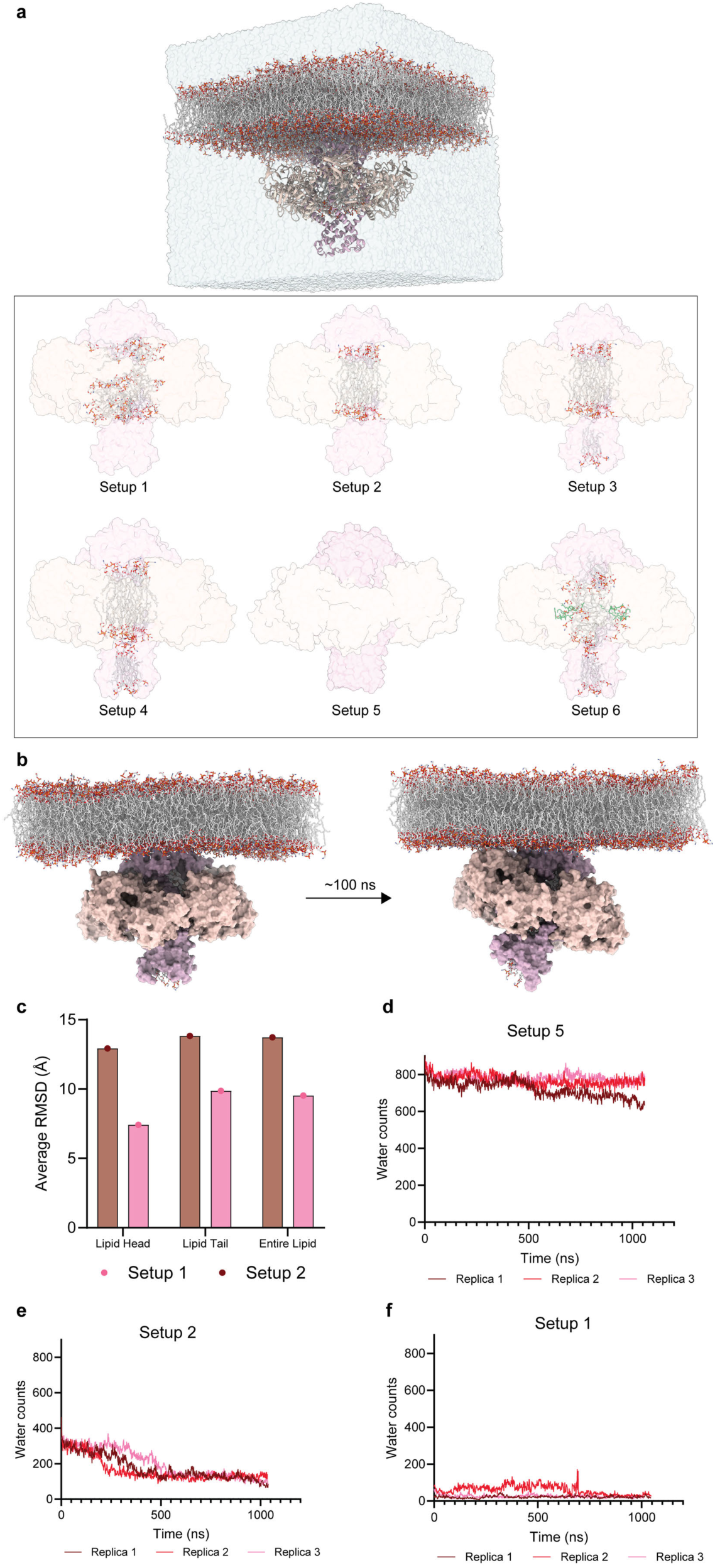
Molecular dynamic simulation setups. a, Model systems constructed in this work. Bottom inset shows the initial lipid arrangement inside the Ncp tube for every simulation setup (**Table S3**), showing the lipids (white) and MK-7 (green). **b,** Tilting of the Ndh-Ncp complex over 100 ns of the simulation trajectory. **c,** The average RMSD (Å) of lipid movement across the simulations in setup 1 and 2, showing lipids are more mobile in setup 2. **d-f,** The number of water molecules within the Ncp chamber over the course of the simulations for setup 5 (no lipids) and setup 2 and 1.

**Supplemental Figure 6.**
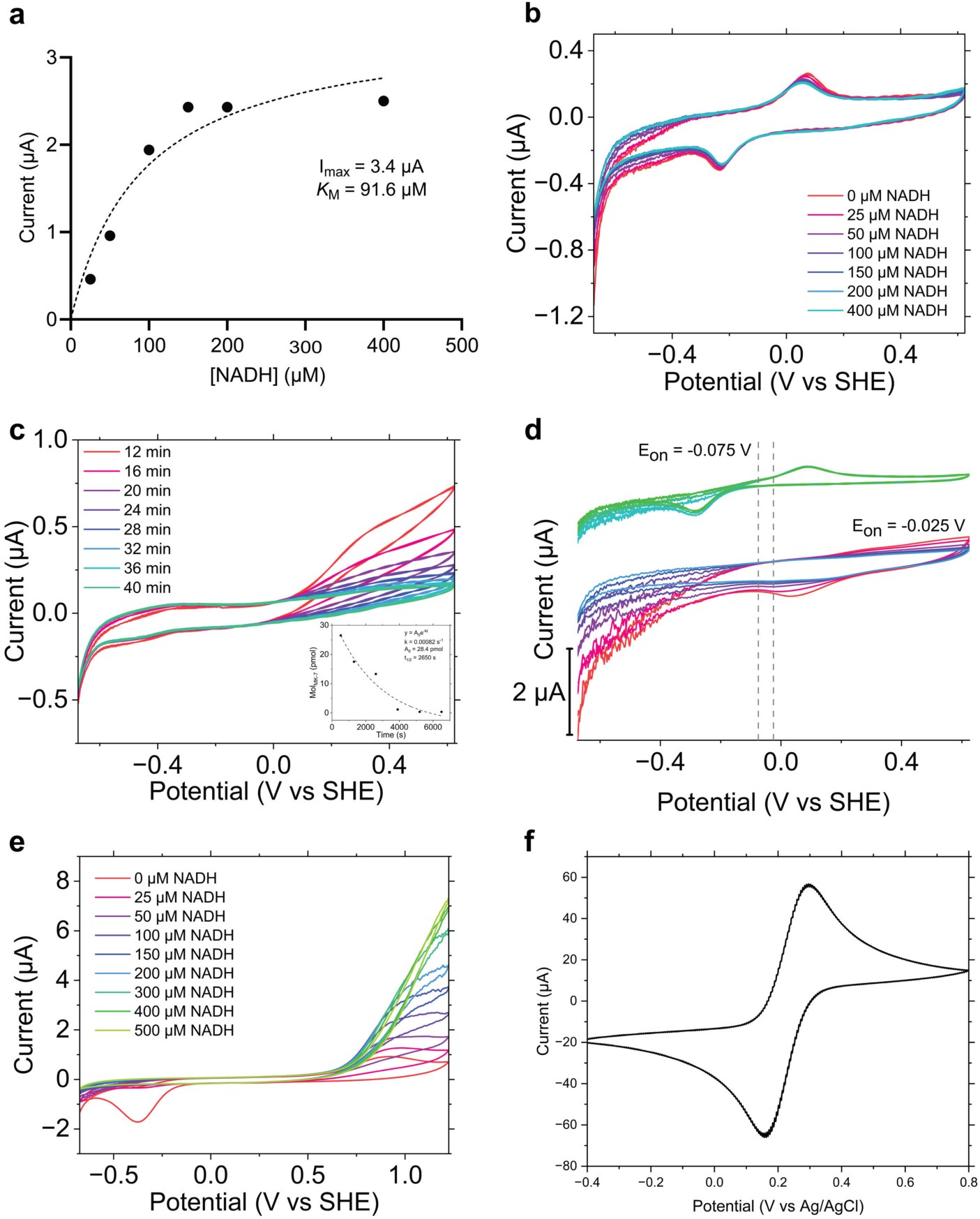
**Cyclic voltammetry of Ndh-Ncp**. **a,** A plot of the current values at +0.3 V vs SHE were extracted from the reductive sweeps (**from Figure 4i**) and plotted against NADH concentration (solid circles), then fit to the Michaelis-Menton equation (dashed line). **b,** Cyclic voltammograms of NADH titration in the absence of Ndh-Ncp, with MK-7 present **c,** Cyclic voltammograms of Ndh-Ncp associated with the bilayer-coated electrode, recorded at 400 μM NADH from-0.675 to +0.625 V vs SHE at 10 mVs^-1^ in the absence of MK-7 at 4-minute intervals, inset of the quinone decay plot calculated from the reductive peak of Figure S5d (bottom). **d,** Cyclic voltammograms of Ndh-Ncp associated with the bilayer-coated electrode in the absence of NADH, in the presence (top) and absence (bottom) of exogenous MK-7. **e,** Cyclic voltammograms recorded at varying NADH concentrations from-0.675 to +1.225 V vs SHE at 10 mVs^-1^ in the presence of ubiquinone-8. **f,** Cyclic voltammogram with a clean, blank electrode recorded in 0.2 M Tris bu_er, pH 7.00 in the presence of 10 mM potassium ferricyanide to calibrate the Ag/AgCl reference electrode against the known midpoint potential of potassium ferricyanide. The scan was recorded from-0.4 to +0.8 V vs Ag/AgCl at 100 mVs^-1^. All data are derived from a single representative experimental, performed after multiple rounds of optimisation.

**Supplemental Figure 7.**
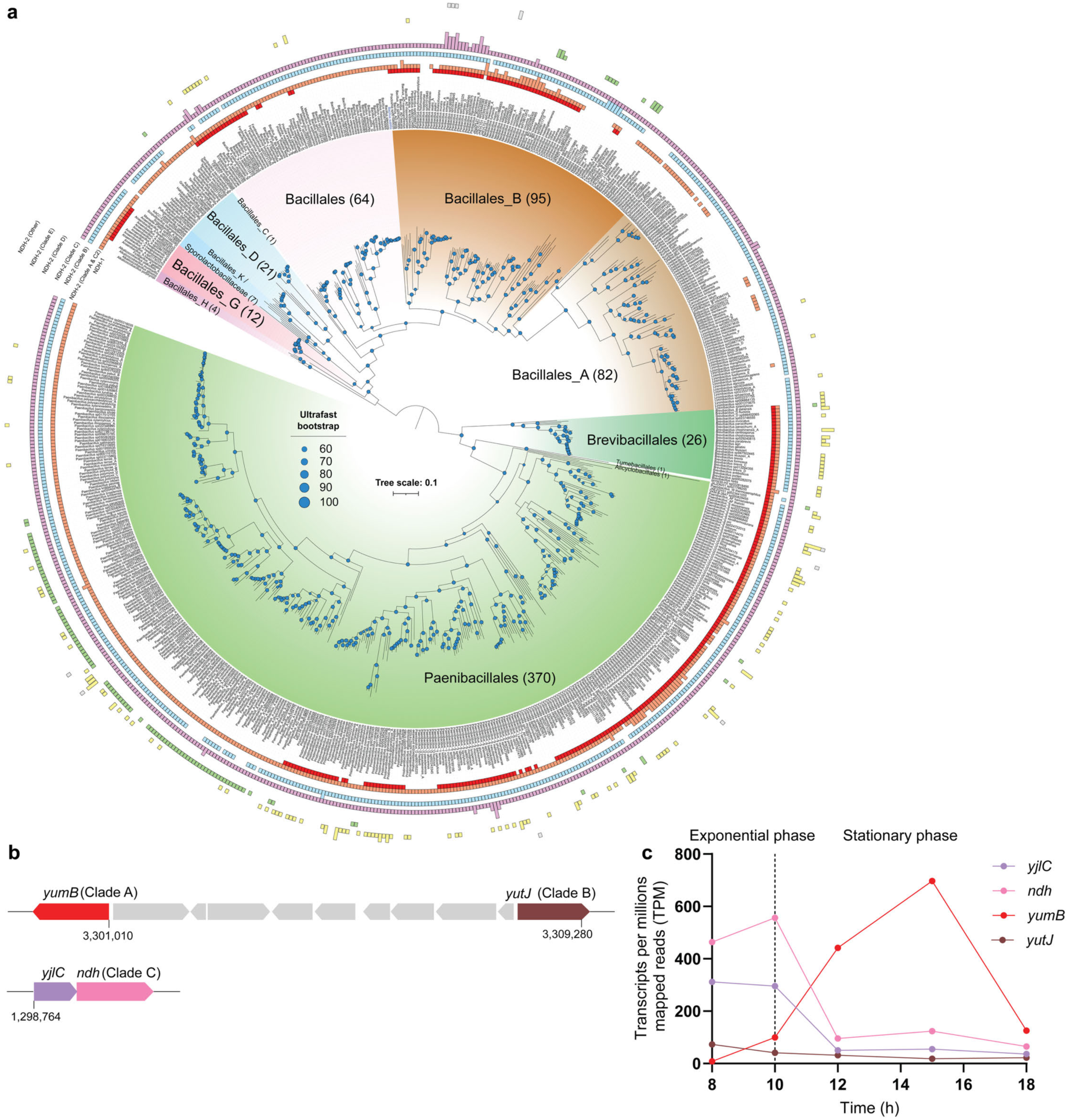
Ndh-Ncp encoding Bacillota are taxonomically widespread and co-encode multiple NDH-2 enzymes with distinct physiological functions. a, Mid-point rooted maximum-likelihood phylogenomic tree of 684 Ndh-Ncp encoding Bacillota species genomes from GTDB R09-RS220 using concatenated 120 GTDB bacterial marker genes and WAG + G20 model. Outer-rings show the copy numbers of Complex I (NDH-1) and various NDH-2 clades encoded by each species. Branch circles show ultrafast bootstrap support values with 1000 replicates. The scale bar corresponds to the expected number of substitutions per site. **b,** Canonical genetic arrangement of the NDH-2 from clade A (*yumB*) and NDH-2 from clade B (*yutJ*), compared to NDH-2 from clade C (*ndh*) and Ncp, using *B. subtilis* as an example. Canonical NADH-oxidizing NDH-2 (Clade A) and NADPH-oxidizing NDH-2 (Clade B) are commonly co-encoded in close genetic proximity (typically 4 – 20 genes apart) in Bacillota genomes. Clade C1 NDH-2 is always co-encoded with Ncp in the immediate genetic proximity **c,** Time-course transcriptomic profiling of NDH-2 and Ncp expression in *B. subtilis* grown in lysogeny broth, from Jun *et al*^52^.

**Supplemental Figure 8.**
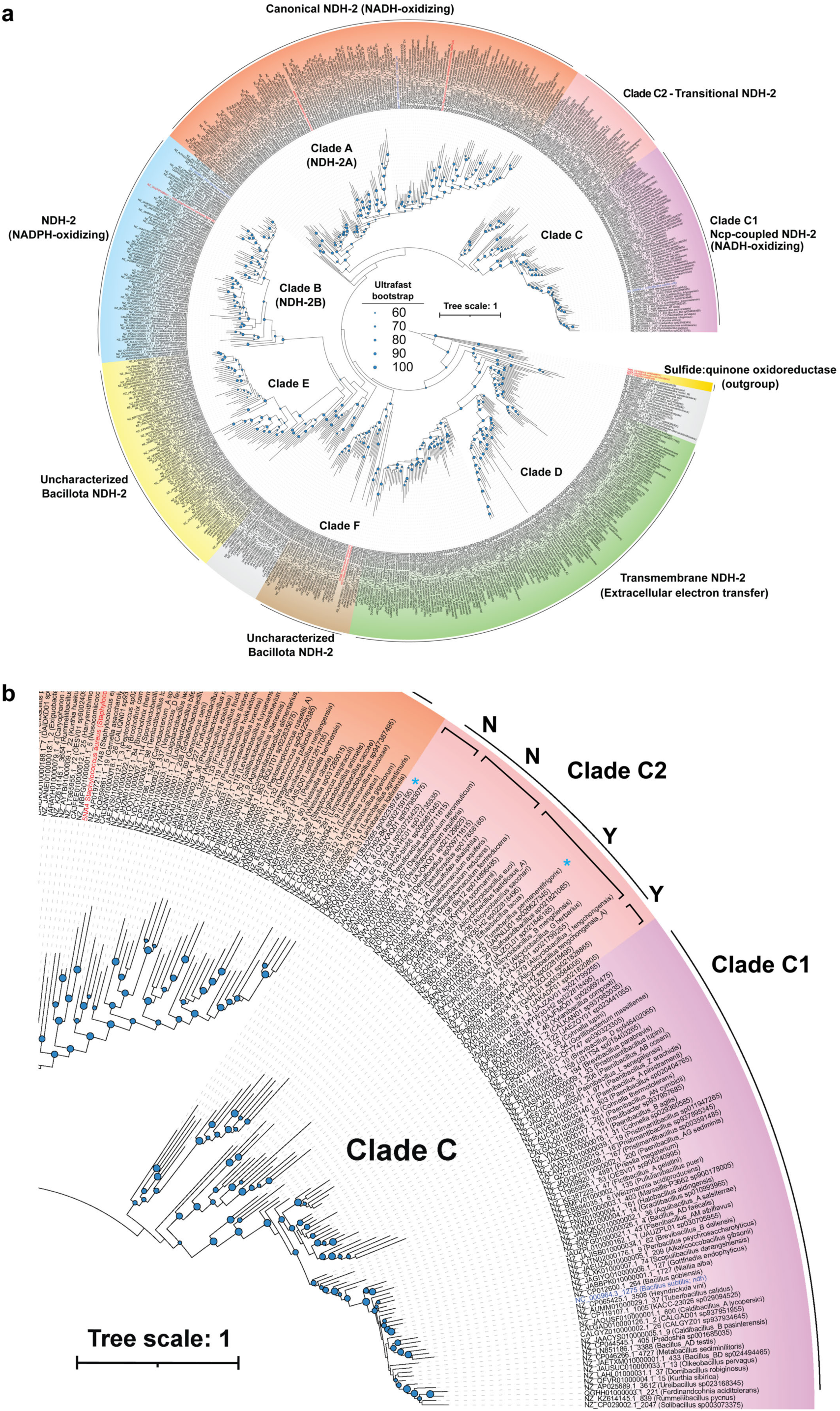
Expanded phylogenetic tree of NDH-2 across Bacillota. a, Maximum-likelihood tree constructed based on 683 cluster-dereplicated NDH-2 amino acid sequences from Bacillota (Ncp-coupled Ndh: 80% identity; Other NDH-2: 60% identity) using the LG+I+R10 model and rooted using sulfide:quinone oxidoreductases (Sqr) as an outgroup. Taxonomy of the genomes follows GTDB R09-RS220. Coloured tip labels show reference NDH-2/Sqr sequences (red) and the three Bacillus subtilis NDH-2 (blue). Branch circles show ultrafast bootstrap support values with 1000 replicates. The scale bar corresponds to the expected number of substitutions per site. **b,** zoom of clade C NDH-2 with annotations on clade C2 showing Ndh with a DUF1641-domain protein present elsewhere in the genome (Y) or absent (N). Those annotated with a blue asterisk are outliers.

**Supplemental Figure 9.**
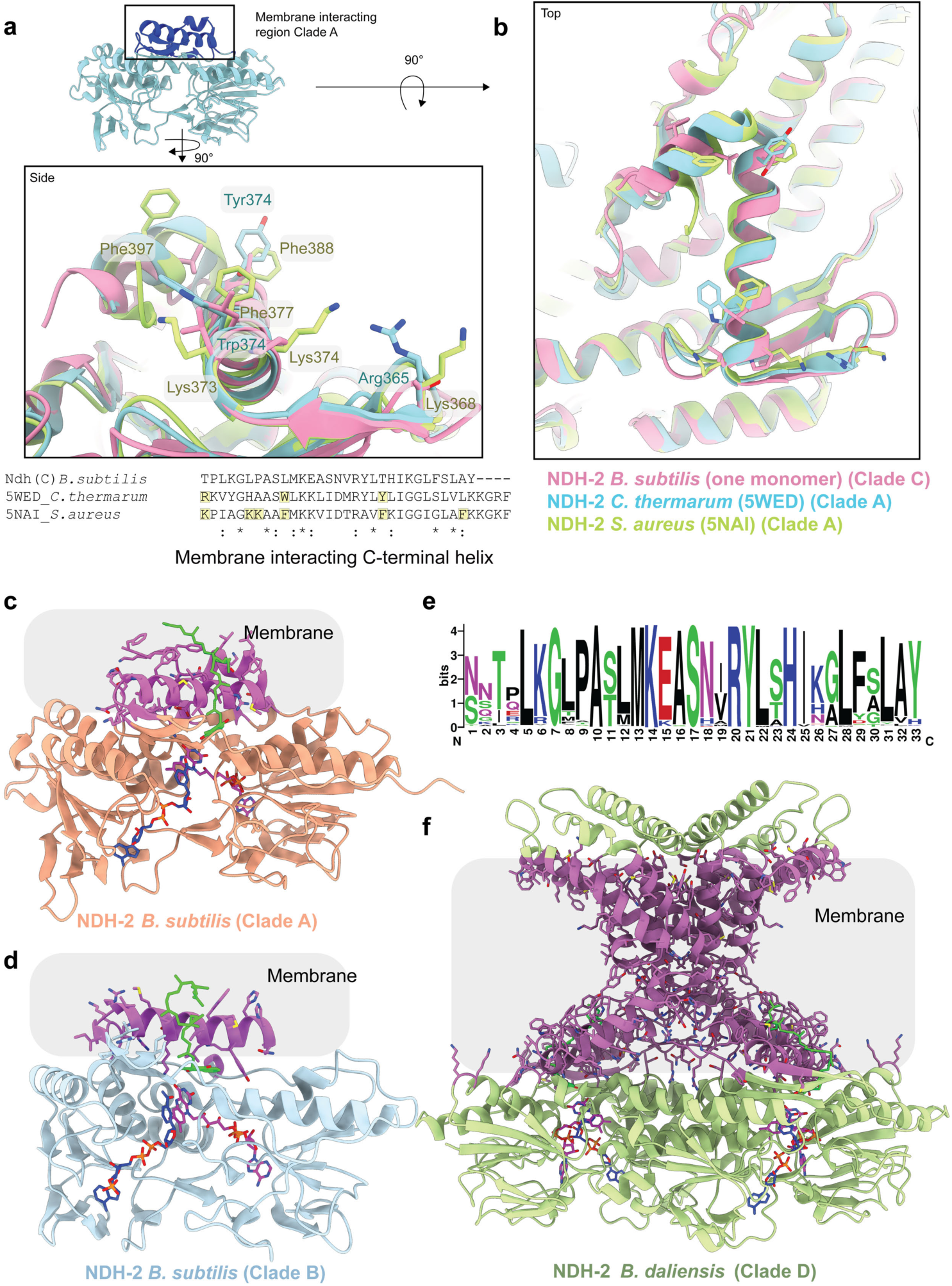
Membrane interaction modes of NDH-2 from Bacillota. a, Overlay of NDH-2 from *B. subtilis* (Clade C1 Ndh, this study), *C. thermarum* (Clade A, PDB: 5WED) and *S. aureus* (Clade A, PDB: 5NAI). The side view of the key residues for membrane interaction from these helices are shown. *S. aureus* (green) and *C. thermarum* (blue) have key aromatic residues that are absent in *B. subtilis* Ndh (pink). **b,** The top view of the structures shows the key residues, with the sequence alignment below demonstrating the conservation of these residues in the clade A NDH-2. **c,** Model of clade A NDH-2 (*B. subtilis*) with membrane-interacting region shown in purple **d,** Model of clade B NDH-2 (*B. subtilis*) **e,** WebLogo of the C-terminus of the Ndh from clade C1 that couple to Ncp from Bacillota **f,** Model showing the membrane-interacting region of clade D NDH-2 (*B. daliensis*). Notably, clade D NDH-2 has transmembrane helices to completely integrate into the membrane.

**Supplemental Figure 10.**
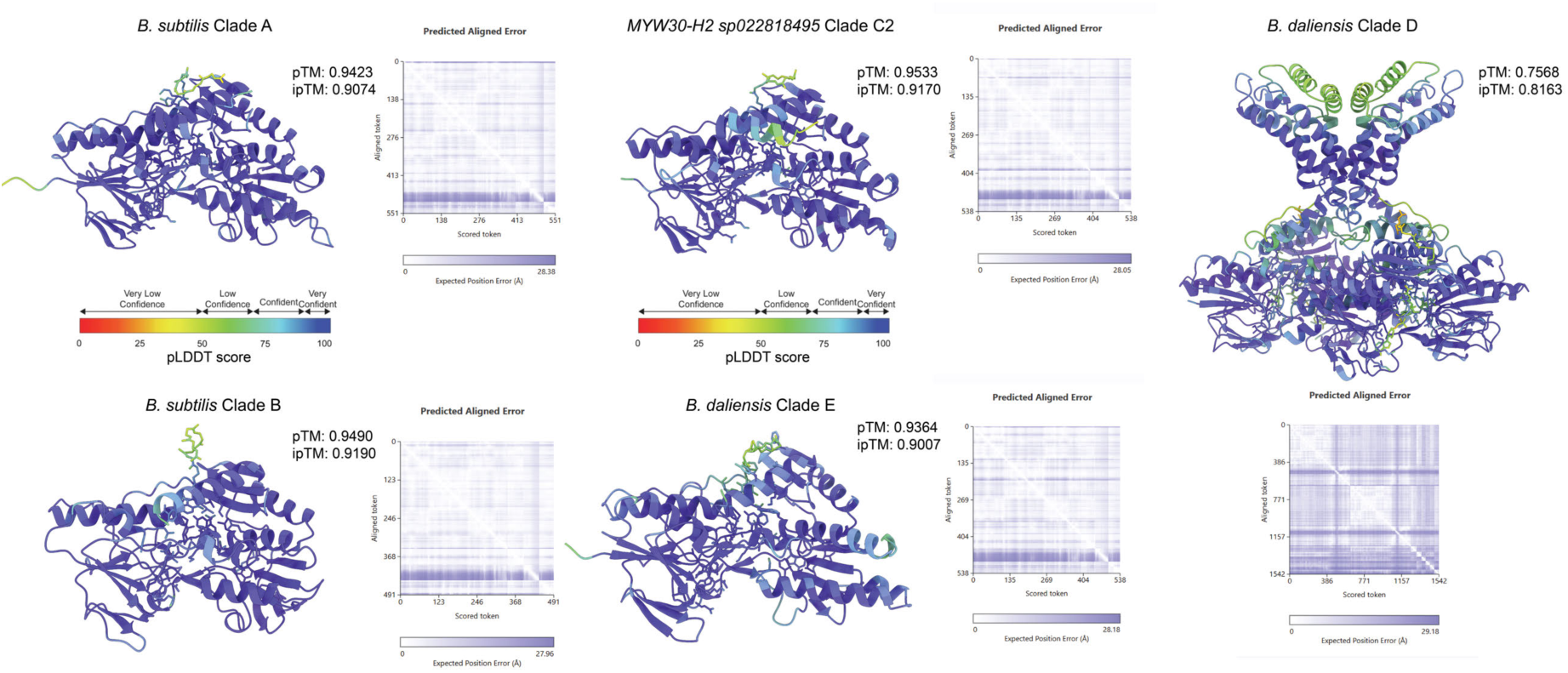
Quality estimates for the Chai1 models of NDH-2 enzymes. NDH-2 enzymes modelled in Figure 5 shown here with colour coding to demonstrate model confidence, indicated with the pLDDT key showing confidence levels. Predicted alignment error (PAE) plots are shown on the right of each model, demonstrating the predicted error of the model, with purple showing less error and white higher error.

**Supplemental Figure 11.**
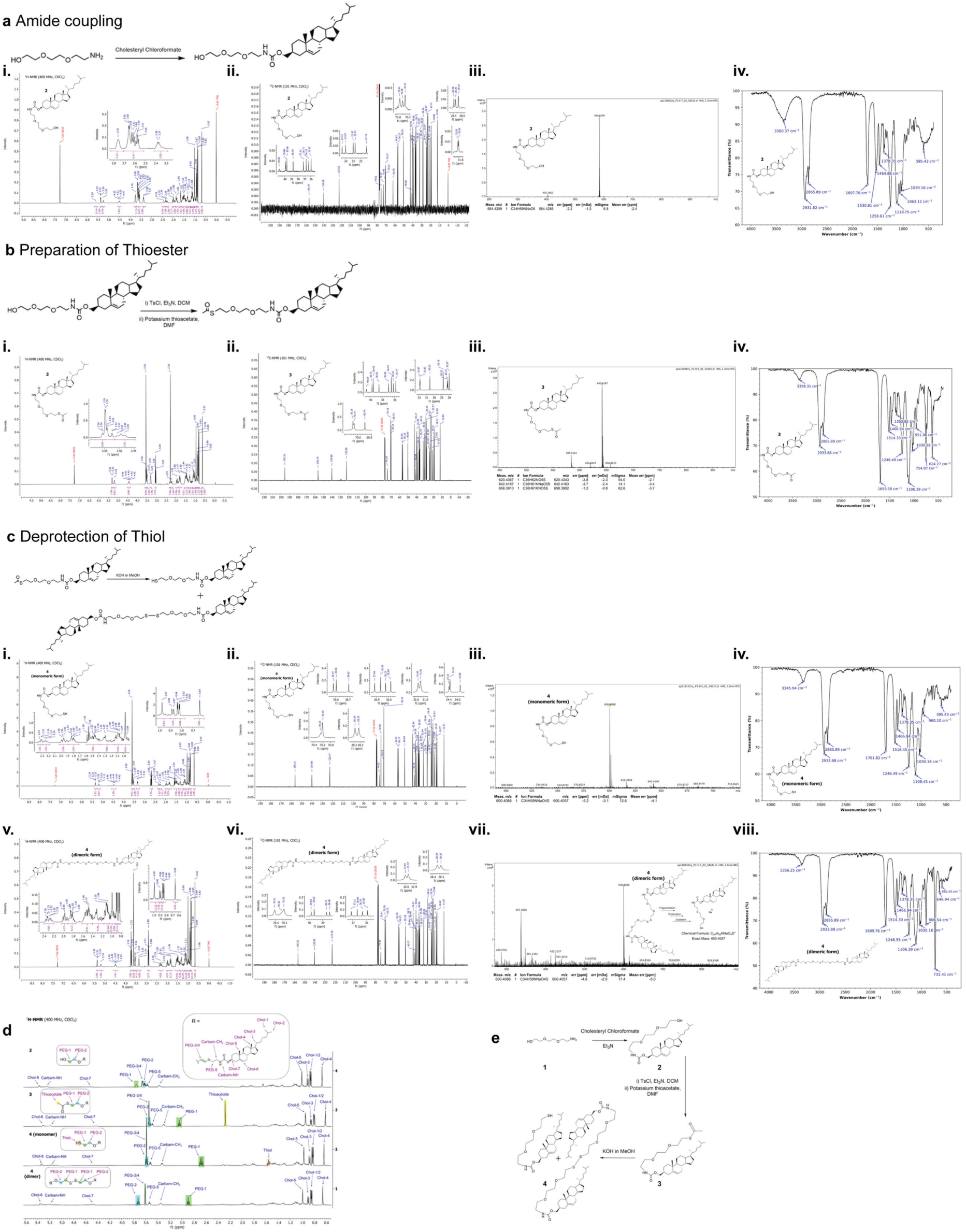
Synthesis and characterisation of S-cholesterol for electrode bilayer. a, Amide coupling reaction. **b,** Preparation of Thioester. **c,** Deprotection of Thiol reaction. **d,** Assignment of readily-distinguishable regions of the ^1^H NMR spectra of compounds 2-4. **e,** The full synthesis of EO-3 cholesterol. All **i.** ^1^H NMR spectrum of each compound, all **ii.** ^13^C NMR spectrum of each compound, all **iii.** (ESI)HRMS spectrum of each compound, all **iv.** FT-IR (ATR) spectrum of each compound. **cv-viii.** is dimeric form of compound 4.

## Supplemental Tables

**Supplemental Table S1.** Cryo-EM data collection, refinement and validation statistics.

|  | *Ndh-Ncp* | *Ndh-Ncp*<br>*filament* | *Ndh-Ncp*<br>*NADH/MK-8* |
| --- | --- | --- | --- |
| **Data collection and processing** |  |  |  |
| Magnification (xk) | 130 | 130 | 96 |
| Voltage (kV) | 300 | 300 | 300 |
| Electron exposure (e-/Å<sup>2</sup>) | 39 | 39 | 50 |
| Defocus range (μm) | -1.2-0.6 | -1.2-0.6 | -1.2-0.6 |
| Pixel size (Å) | 0.655 | 0.655 | 0.808 |
| Symmetry imposed | C4 | C4 | C4 |
| Initial particle images (no.) | 483,556 | 483,556 | 5,095,952 |
| Final particle images (no.) | 356,549 | 389,955 | 341,297 |
| Map resolution (Å) | 2.11 | 3.09 | 2.52 |
| FSC threshold | 0.143 | 0.143 | 0.143 |
| Map resolution range (Å) | 2.1-3.1 | 2.6-4.6 | 2.1-4.3 |
| **Refinement** |  |  |  |
| Initial model used (PDB code) | - | - | - |
| Model resolution (Å) | 2.11 | 3.09 | 2.52 |
| FSC threshold | 0.143 | 0.143 | 0.143 |
| Model resolution range (Å) | n/a | n/a | n/a |
| Map sharpening *B* factor (Å<sup>2</sup>) |  |  |  |
| Model composition |  |  |  |
| Non-hydrogen atoms | 15,768 | 46,296 | 16,084 |
| Protein residues | 2,044 | 6,004 | 2,044 |
| Ligands | 4 | 12 | 16 |
| *B* factors (Å<sup>2</sup>) |  |  |  |
| Protein | 87 | 156 | 95 |
| Ligand | 65 | 105 | 103 |
| R.m.s. deviations |  |  |  |
| Bond lengths (Å) | 0.002 | 0.003 | 0.002 |
| Bond angles (°) | 0.486 | 0.538 | 0.514 |
| Validation |  |  |  |
| MolProbity score | 1.16 | 1.38 | 1.31 |
| Clashscore | 3.71 | 5.50 | 5.66 |
| Poor rotamers (%) | 0 | 0.00 | 0 |
| Ramachandran plot |  |  |  |
| Favored (%) | 98.57 | 97.63 | 98.72 |
| Allowed (%) | 1.28 | 2.37 | 1.28 |
| Disallowed (%) | 0.15 | 0.00 | 0.00 |

**Supplemental Table S2.**
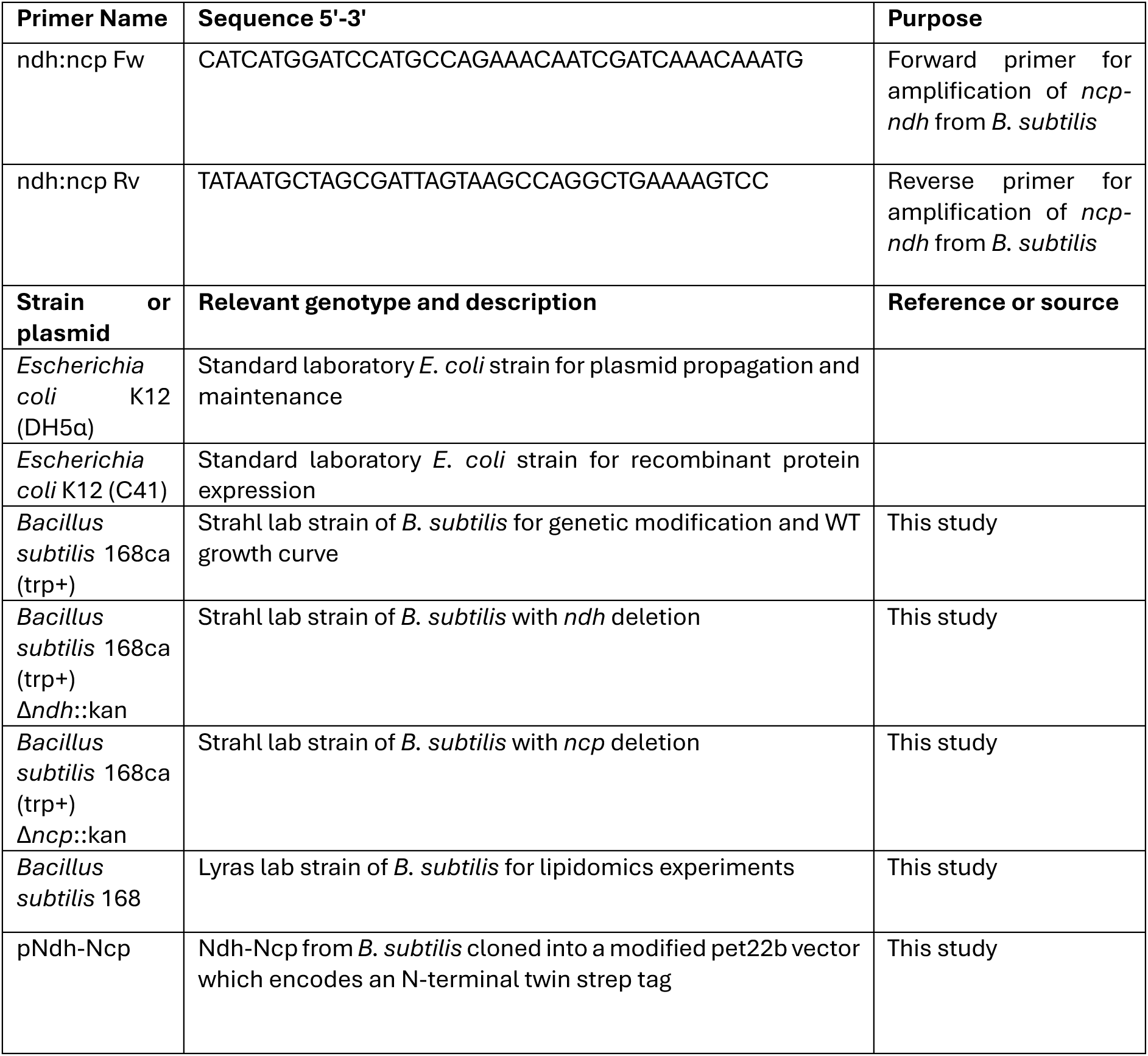
Primers, strains and plasmids used in this study.

**Supplemental Table S3.**
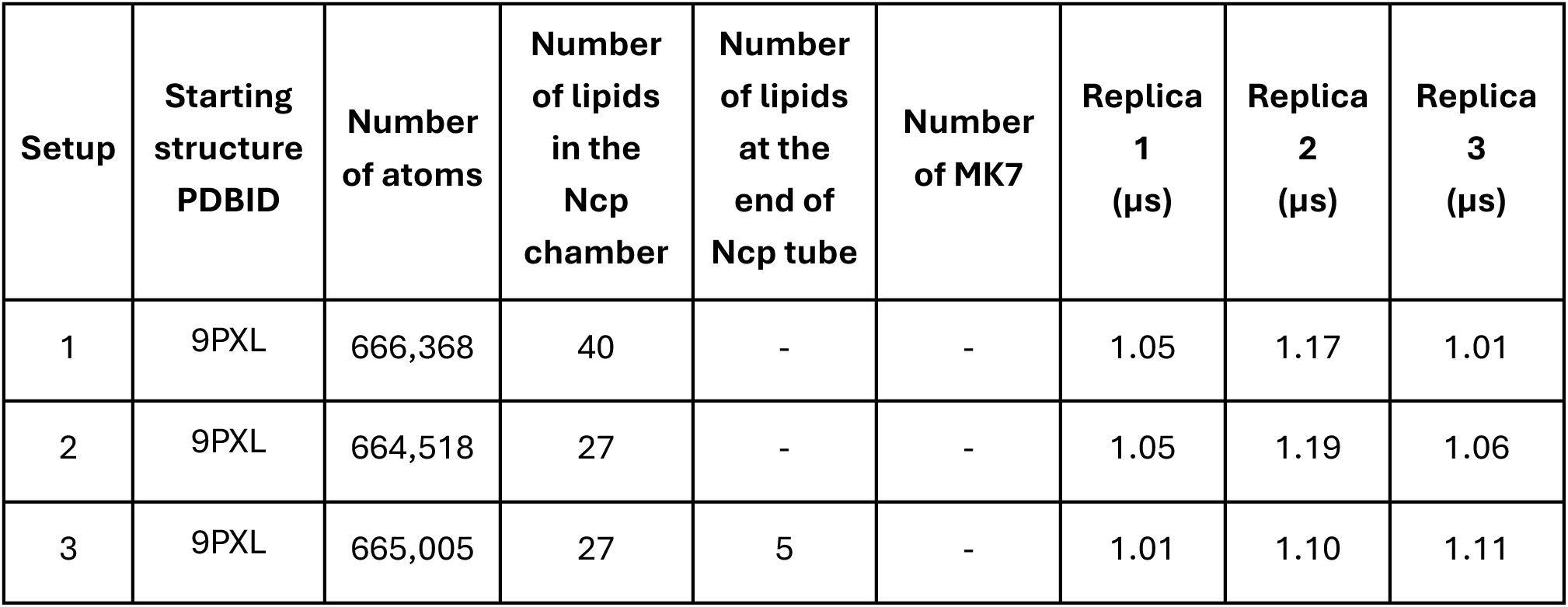
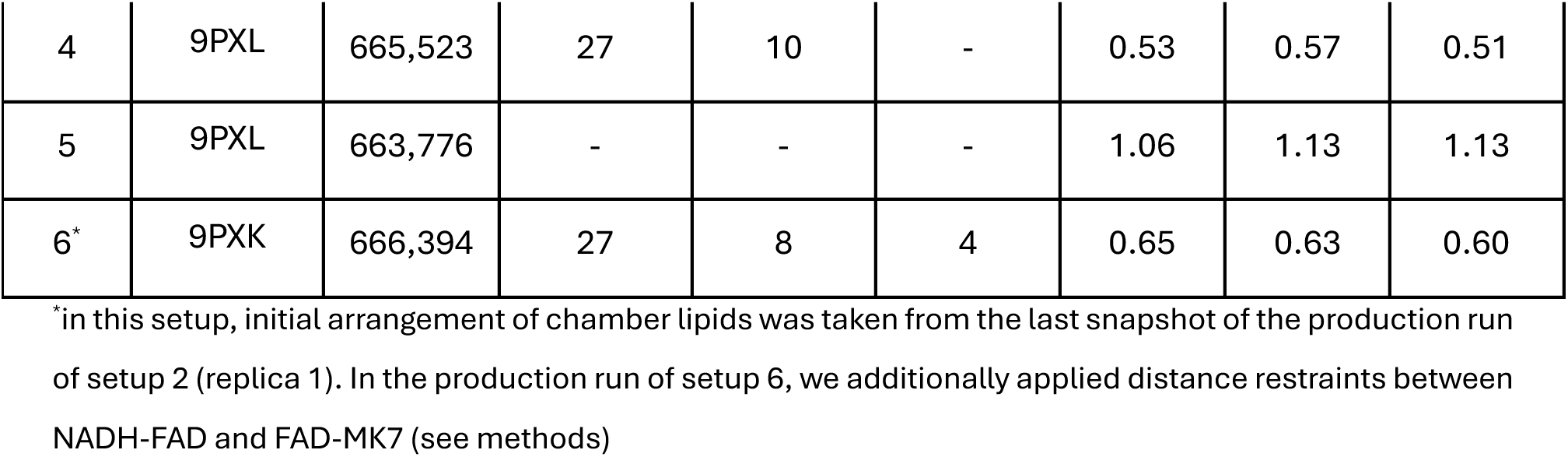
Classical MD simulation setups, their size, time scales, and lipid composition.

